# Nrd1p identifies aberrant and natural exosomal target messages during the nuclear mRNA surveillance in *Saccharomyces cerevisiae*

**DOI:** 10.1101/2020.06.02.129106

**Authors:** Pragyan Singh, Anusha Chaudhuri, Mayukh Banerjea, Neeraja Marathe, Biswadip Das

## Abstract

In all eukaryotes, selective nuclear degradation of aberrant mRNAs by nuclear exosome and its cofactors TRAMP, and CTEXT contribute to the fidelity of the gene expression pipeline. In the model eukaryote, *Saccharomyces cerevisiae*, the Nrd1p-Nab3p-Sen1p (NNS) complex, previously known to be involved in the transcription termination and matured 3’-end formation of vast majority of non-coding and several coding RNAs, is demonstrated to universally participate in the nuclear decay of various kinds of faulty messages in this study. Consistently, *nrd1-1/nrd1-2* mutant cells display impairment of the decay of all kinds of aberrant mRNAs, like the yeast mutants deficient in Rrp41p, Rrp6p, and Rrp4p. *nrd1*Δ^CID^ mutation (consisting of Nrd1p lacking its CID domain thereby abrogating its interaction with RNAPII) however, abolishes the decay of aberrant messages generated during early phases of mRNP biogenesis (transcription elongation, splicing and 3’-end maturation) without affecting the decay rate of the export-defective mRNAs. Mutation in the 3’-end processing factor, Pcf11p, in contrast, displayed a selective abolition of the decay of the aberrant mRNAs, generated at the late phase of mRNP biogenesis (export-defective mRNAs) without influencing the faulty messages spawned in the early phase of mRNP biogenesis. Co-transcriptional recruitment of Nrd1p onto the faulty messages, which relies on RNAPII during transcription elongation and on Pcf11p post transcription, is vital for the exosomal decay of aberrant mRNAs, as Nrd1p deposition on the export-defective messages led to the Rrp6p recruitment and eventually, their decay. Thus, presence of the ‘Nrd1p mark’ on aberrant mRNAs appears rate-limiting for the distinction of the aberrant messages from their normal functional counterparts.

**Author’s Summary:** Aberrant/faulty mRNAs generated from the deficiencies in any of the mRNP biogenesis events are promptly eliminated by the nuclear exosome and its cofactors TRAMP and CTEXT complexes. These machineries work relentlessly in the nucleus to detect all kinds of aberrant mRNAs and selectively target them for destruction. However, initial detection of a minuscule amount of aberrant mRNA in the vast background of normal mRNAs is quite challenging and its mechanism remains elusive. In this work, we demonstrate that, the trimeric Nrd1p-Nab3p-Sen1p complex, previously implicated in the transcription termination of diverse non-coding RNAs and a handful of mRNAs, constitute an integral component of the nuclear mRNA surveillance mechanism in baker’s yeast *Saccharomyces cerevisiae*. Major component of this complex, Nrd1p is demonstrated to be recruited selectively onto various classes of representative model aberrant messages either co-transcriptionally by RNA Polymerase II or post-transcriptionally by Pcf11p. Binding of Nrd1p to the export-defective special mRNAs further leads to the recruitment of Rrp6p on to them thereby leading to their degradation. NNS complex thus plays a vital role of initially recognizing the faulty messages and further assists in the recruitment of the nuclear exosome for their prompt elimination.

## Introduction

Aberrant messages derived from the inaccurate mRNP biogenesis are eliminated by a broad spectrum of mRNA surveillance and quality control mechanisms (1–4). In *Saccharomyces cerevisiae*, the nuclear exosome, along with its cofactors, degrade a wide array of aberrant and normal mRNAs (1,2,5–8). Functionally, these cofactors recognize specific RNA targets and then further recruit them to the core exosome (EXO11^Dis3p+Rrp6p^) to promote their degradation (9). TRAMP (**<u>TR</u>**f4p/5p-**<u>A</u>**ir1p/2p-**<u>M</u>**tr4p-**<u>P</u>**olyadenylation) complex, the best-studied cofactor in *S. cerevisiae* consists of DExH box RNA helicase, Mtr4p (4,10,11), non-canonical poly(A) polymerase, Trf4p/Trf5p and Zn-knuckle RNA binding proteins, Air1p/2p (4,10,11). In addition to TRAMP, two other nuclear cofactors, CTEXT (**<u>C</u>**bc1p-**<u>T</u>**if4631p-dependent **<u>EX</u>**osomal **<u>T</u>**argeting) (previously termed as DRN) (12–16), and NNS (Nrd1p-Nab3p and Sen1p) complexes (17–22) assist the exosome in targeting both aberrant/normal mRNAs as well as a vast majority of ncRNAs (sno-/sn-/CUTS, NUTs and SUTS). CTEXT consists of nuclear cap-binding protein Cbc1p/2p (13,14), shuttling proteins Tif4631p/ Upf3p (16), a DEAD-box RNA helicase, Dbp2p (23) and meiotic protein Red1p (24), and degrades a distinct group of aberrant (12–16) and normal mRNAs (25–27). The NNS complex, in contrast, comprises of Nrd1p and Nab3p as the two major sequence-specific RNA binding proteins (18) and Sen1p as the major DNA-RNA helicase (28) (see below).

Aberrant mRNAs in the yeast nucleus were classified into early, intermediate, and late depending on the specific phases of mRNP biogenesis events at which they are generated (Table 1). The transcription-elongation and splice-defective messages were classified as early, the aberrant 3’-end processing-defective transcripts were categorized as intermediate, and the export-defective messages were classified as the late category (12). Remarkably, while the nuclear exosome degrades all of these messages, the TRAMP degrades only the faulty messages derived in the early phase, and the CTEXT degrades only the faulty transcripts produced during the terminal stage of mRNP biogenesis (12). Strikingly, the degradation of aberrant messages derived at the intermediate stage of mRNP biogenesis requires the involvement of both TRAMP and CTEXT (12). However, the molecular basis of the mRNP-biogenesis-stage-specific participation of TRAMP and CTEXT onto the distinct classes of aberrant messages is still unclear.

**Table 1:**
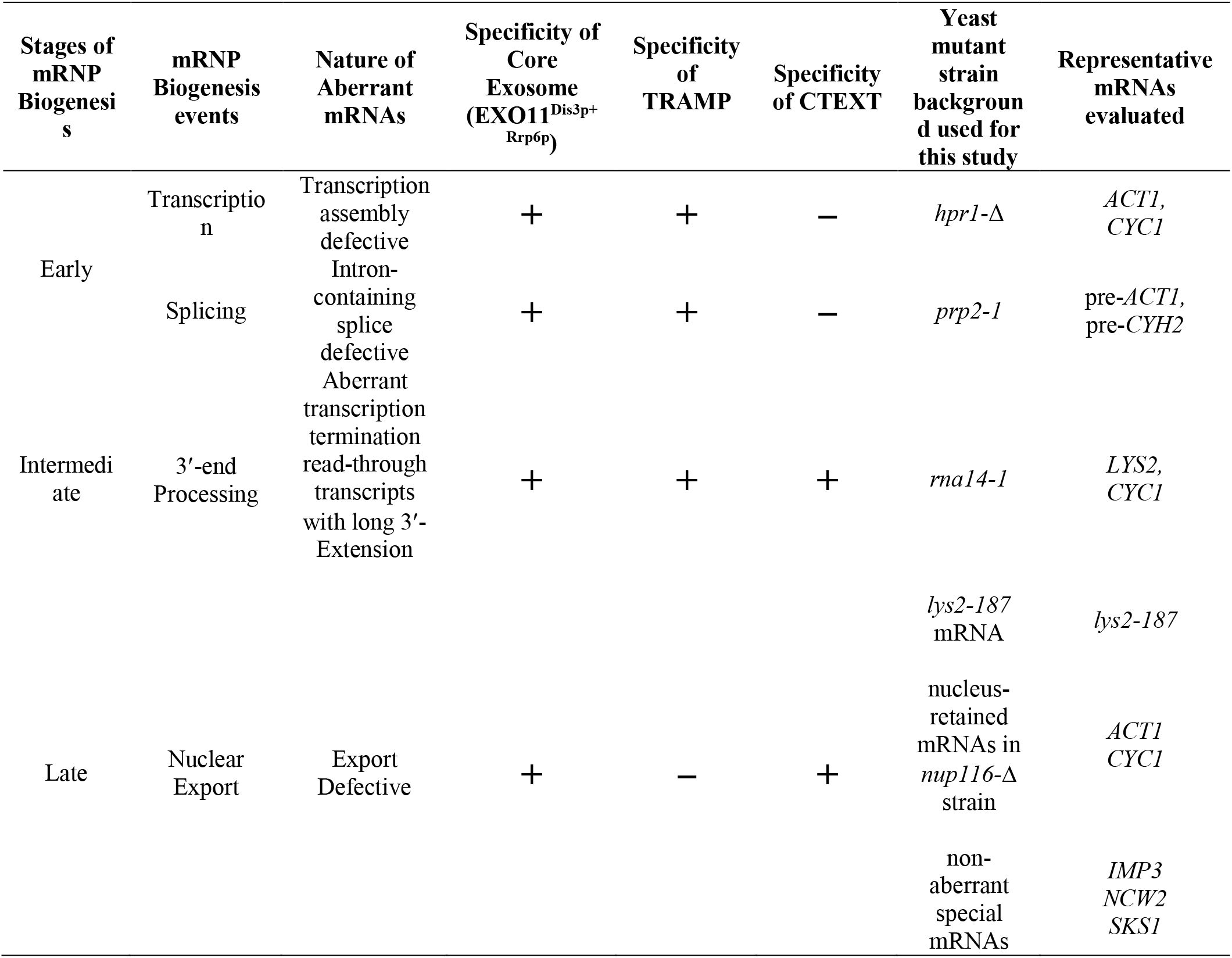
Various types of aberrant mRNA substrates, generated progressively during mRNP biogenesis, the specificity of the decay apparatus to degrade them, and their representative model mRNAs used in this study.

In *Saccharomyces cerevisiae*, an alternative pathway of transcription termination exists besides the canonical pathway, which influences the stability of a few protein-coding transcripts (21,29). The RNA binding protein Nrd1p, complexed with its partners, Nab3p and Sen1p (a putative helicase), dubbed NNS (Nrd1p-Nab3p-Sen1p) complex, plays a pivotal role in transcription termination of pre-snRNAs, pre-snoRNAs, and Cryptic Unstable Transcripts (CUTs) in this alternative mechanism (17–19, 22, 30–32), thereby leading either to their maturation or degradation by the exosome and TRAMP (20,33). This trimeric complex was demonstrated to bind their RNA targets via the interactions through the elongating RNA Polymerase II (RNAPII) as well as the nascent RNA (18). Typically, the serine 2 (Ser-2) and 5 (Ser-5) residues of the repetitive C-terminal domain of RNAPII predominantly undergo reversible phosphorylation at various phases of transcription elongation and thereby influence various events of mRNP biogenesis via the recruitment of diverse protein factors involved in these mRNA processing events (34–37). Other than Ser-2 and 5, Tyr-1, Thr-4, and Ser-7 also contribute to the co-transcriptional recruitment of the various processing factors by undergoing reversible phosphorylation (21,38). In the early elongation phase, the Ser-5 and Tyr-1 phosphorylation of the CTD of RNAPII predominates (up to about 100 nucleotides for the yeast genes). After about 100 nucleotides, the phosphorylated ser-5 starts to decrease, and phosphorylated ser-2 starts to increase (38). Although Nrd1 binding is reported to be favored by Ser-5 and Tyr-1 phosphorylation (39,40), the role of Tyr-1 phosphorylation in this process is debatable since several studies have demonstrated that Tyr-1 phosphorylation, which remains very low near the transcription start site (TSS), strongly antagonizes Nrd1p binding (21,38). Nevertheless, binding of Nrd1p and Nab3p to the nascent mRNAs towards the TSS of the protein-coding mRNA (38) subsequently recruits Sen1p helicase, which promotes dissociation of the RNAPII possibly by destabilizing the RNA-DNA hybrid in the transcription bubble (28,39,41,42).

NNS complex directs the decay of antisense mRNA/transcripts, and several protein-coding mRNAs (43–47) that includes the *NRD1* mRNA (to control its expression) (17,19), *URA8, URA2, ADE12* (48,49) and *FKS2* (50). Interestingly, the binding motif to which Nrd1 and its binding partner Nab3p prefer to bind is poorly represented in mRNAs and is highly enriched in sn- and snoRNAs (22). However, Nrd1p and Nab3p were shown to bind to the hundreds of protein-coding mRNAs (22,43–45,51,52), the functional importance of which remained unclear. Although a couple of studies demonstrated a correlation between Nrd1p/Nab3p/Mtr4p binding to stress-responsive messages during glucose starvation with their decay (43,51), the exact nature of the functional role of NNS complex in the decay of these mRNAs remained elusive.

In addition to the normal and functional mRNAs, the NNS complex was also shown to direct the degradation of bacterial Rho factor-induced aberrant transcripts that involved the Nrd1p-dependent coordinated recruitment of Rrp6p after being recruited by the RNAP II (46). Genome-wide high-resolution landscapes of Rrp6p, Trf4p, and Nrd1p/Nab3p were utilized to show that under normal conditions, Nrd1p/Nab3p appeared to withdraw from mRNA loci and sequester around the sno- and snRNA loci in the genome. Upon activation of the Rho factor that induces the formation of aberrant mRNP lacking the appropriate complement mRNA maturing factors, Nrd1p/Nab3p promptly redistribute from the genomic loci producing non-coding RNAs to the new loci harboring Rho-affected protein-coding genes, thereby triggering their decay and elimination (47). However, whether Nrd1p complex universally participates in the degradation of all kinds of aberrant messages and via what mechanism it carries out the surveillance has never been studied. In this investigation, we present evidence that Nrd1p (and presumably the NNS complex) plays a central role in the surveillance of the entire spectrum of aberrant nuclear mRNAs. Moreover, the co-transcriptional recruitment of Nrd1p (i) on all kinds of faulty transcripts is found to be crucial for their decay, and (ii) on the export-defective mRNAs leads to the recruitment of the exosome component Rrp6p. Our evidence suggests that mode of recruitment of Nrd1p onto a given aberrant message is also vital to govern if it would further facilitate the TRAMP- or CTEXT-dependent degradation of a distinct class of faulty messages.

## Results

We initiated this study with a major pursuit (i) if the NNS complex plays a universal role in the nuclear degradation of the entire spectrum of aberrant messages, and (ii) to unfold the functional relationship of NNS with the TRAMP/CTEXT/exosome. Using a holistic approach, we analyzed its functional role in the nuclear decay of various kinds of aberrant messages (Table 1) by comparing the steady-state levels and stability of arbitrarily chosen representative model mRNAs from different aberrant classes in isogenic series of strains (Fig 1A). These series consist of a parent strain defective in a given type of mRNP biogenesis and its isogenic derivative, which additionally harbor various mutations in the *NRD1* gene (Fig. 1A). The stability analysis was then followed by the systematic investigations of Nrd1p and Rrp6p occupancy profiles onto various kinds of aberrant messages to gain an insight into the mechanism of how the Nrd1p recruitment is taking place onto a specific type of aberrant message and how its recruitment further influence the downstream fate of the aberrant messages.

**Figure 1.**
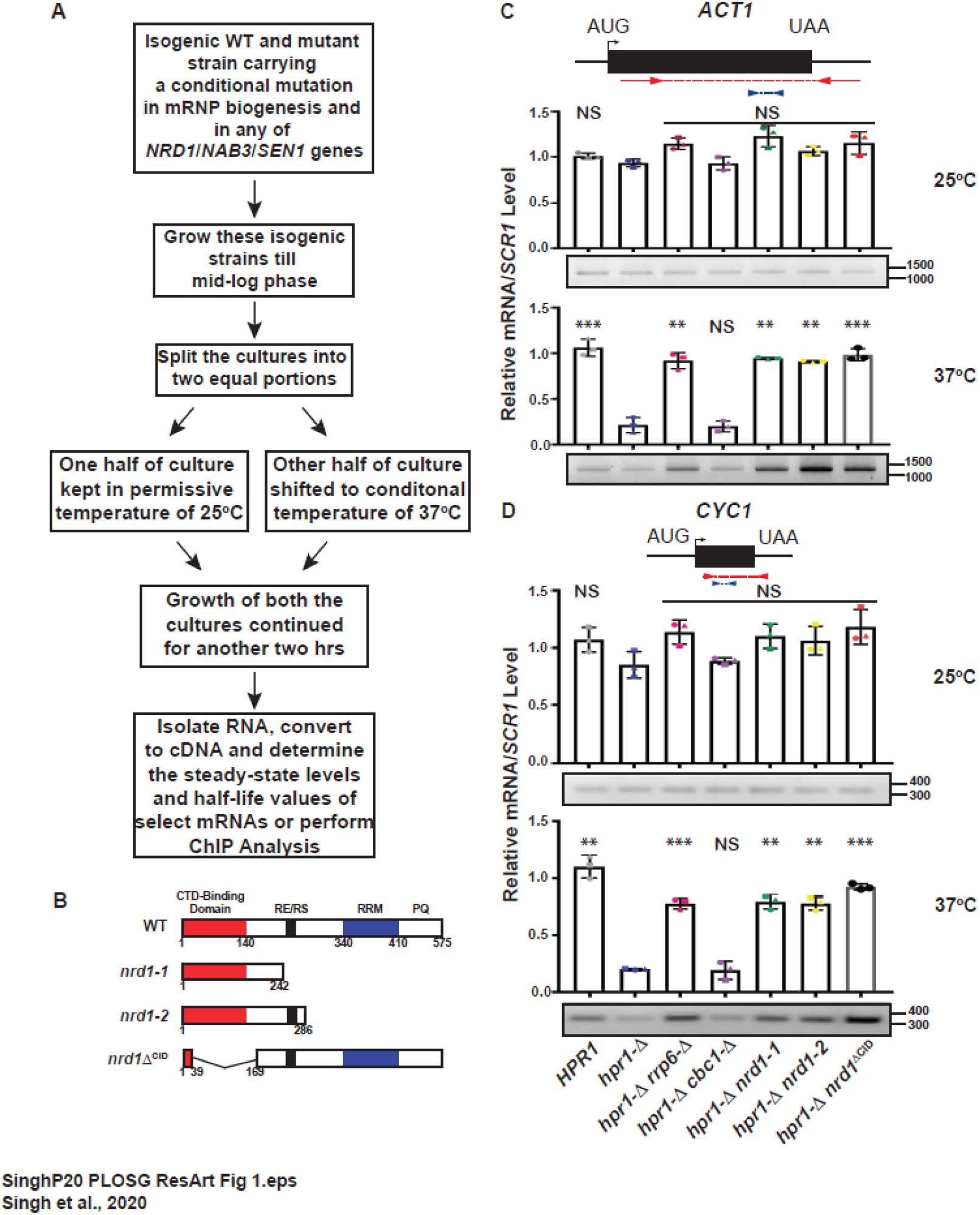
Nrd1p participates in the degradation of aberrant mRNPs generated in transcription-elongation-defective *hpr1*-Δ yeast strain. (A) Outline of the experimental procedure used in this investigation. (B) Schematic diagram showing the domain organization of full-length Nrd1p (redrawn from reference number (91)): the RNAPII CTD binding domain (CID, aa 1–140), RNA Recognition Motif (RRM, aa 340–410), RE/RS (arginine-, serine-, and glutamate-rich region, aa 245–265); and P/Q (proline-, glutamine-rich region, aa 500–575) regions. The specific structures for the *nrd1* mutants are shown below. (C-D) Relative abundance of transcription assembly-defective *ACT1* (panels C) and *CYC1* (panels D) mRNAs at permissive (25°C) and non-permissive temperature (37°C) in the *HPR1^+^, hpr1*-Δ, and its various isogenic derivative strains as determined by qRT-PCR using a primer set encompassing the amplicons indicated in blue on top. The mean normalized level of each target mRNA estimated in the *HPR1*^+^ sample was set to 1. P-value was estimated with respect to the steady-state levels of the transcript in the *hpr1*-Δ yeast strain. The gel panels below each of the graph depicts a representative gel showing the relative levels of the nearly full-length messages as determined by the end-point PCR carried out using a primer set encompassing the amplicon indicated in red on top. Positions of the relevant size standards are indicated at the right side of each gel panel. Three independent samples of random-primed cDNA (from biological replicates, N=3) were prepared from indicated strains, pre-grown at 25°C followed by a 2-h shift at 37°C before subjected to either qRT-PCR or end-point PCR analysis. Transcript copy numbers/3 ng cDNA were normalized to *SCR1* RNA signals obtained from respective samples and are shown as means ± SE. P values were estimated from Student’s two-tailed t-test for a given pair of test strains for each message and is presented with the following symbols, * <0.05, **<0.005 and ***<0.001, NS, not significant.

### Nrd1p participates in the degradation of aberrant mRNAs generated in the early, intermediate and late phases of mRNP biogenesis

During transcription elongation of the protein-coding genes in eukaryotes, a complex of Hpr1p, Tho2p, Mft1, Thp2 (dubbed THO complex) is deposited onto transcribing messages, which leads to its subsequent association with the splicing/export factors Sub2p/Yra1p to promote their mRNP maturation and export (53–55). Mutation in any of the THO components (generating a *hpr1*-Δ or *mft1*-Δ strain) led to the destabilization of several transcripts, (53,55,56), which were rescued by the inactivation of the nuclear exosome, and TRAMP, but not by the inactivation of CTEXT (12). Similarly, a mutation in the *PRP2* gene (generating a *prp2-1* strain) encoding an essential splicing factor (57) resulted in the accumulation of splice-defective pre-mRNAs in the nucleus at a non-permissive temperature of 37°C, which were rapidly degraded by the nuclear exosome (58) and TRAMP complex (12,59). In the conditionally lethal *rna14-1/rna15-2* strains, defective in 3’-end processing, globally defective transcripts are readily produced at restrictive temperature of 37°C, which harbor aberrantly long 3’-extended read-through termini (60–63). The nuclear exosome processes these defective transcripts to their natural length (53,64), and consequently, they were found to be stabilized by the mutations in the components of either the nuclear exosome, TRAMP, or CTEXT (12). These data collectively suggested that assembly- and splice-defective mRNPs in *hpr1*-Δ and *prp2-1* strains (early) undergo degradation in the exosome/TRAMP-dependent manner. In contrast, the decay of the read-through transcripts in *rna14-1*/*15-2* strains (intermediate) requires the exosome/TRAMP/CTEXT complex and the export-defective mRNAs (late) require the action of the exosome/CTEXT (Table 1) (12).

To assess the role of Nrd1p in the decay of the transcription-elongation-, splice- and 3’-end processing-defective transcripts, we initially determined the steady-state levels of two arbitrarily chosen model mRNAs produced in each of the three isogenic yeast strain series at a non-permissive temperature of 37°C (12). These series consist of a parent strain defective in transcription elongation (*hpr1*-Δ), splicing (*prp2-1*), and 3’-end processing (*rna14-1*) events and derivative of this parent with various mutations in *NRD1* gene as indicated in Table S1, Figs 1 and 2. We used three different mutations in the *NRD1* gene (Fig 1B), the first two (*nrd1-1*, and *nrd1-2*) harbor pre-mature non-sense mutations leading to the formation of two truncated versions of Nrd1p. The last one (*nrd1*Δ^CID^) carries a deletion of its CID (RNAPII **<u>C</u>**TD **<u>I</u>**nteracting **<u>D</u>**omain) that abolishes the ability of the mutant *nrd1*Δ^CID^ protein to interact with RNAPII (20,33) (Fig 1B). Remarkably, all of the test transcripts (THO-depleted aberrantly-packaged *ACT1*, and *CYC1* mRNAs *in hpr1*-Δ background, intron-containing *pre-ACT1* and *pre-CYH2* precursor messages in the *prp2-1* background, and aberrantly long 3’-extended read-through transcripts of *LYS2* and *CYC1* genes in the *rna14-1* background) displayed a robust stabilization (varying from three to nine-folds in different mutant strains) in *nrd1-1*, *nrd1-2*, and *nrd1*Δ^CID^ strains at the non-permissive temperature of 37°C (Figs 1 and 2). It should be noted here that the careful selection of primer-sets and subsequent designing of the amplicons for each model mRNA analyzed in this study either by qRT-PCR (data of which presented as bars in Figs. 1 and 2) or by endpoint PCR (data presented as gels at the bottom of graphs) were also used in our previous analysis (12). Use of these primer sets enabled us to unequivocally distinguish the intron-containing precursor mRNAs of *ACT1* and *CYH2* formed in *prp2-1* strain and the aberrant read-through transcripts of *LYS2* and *CYC1* mRNAs with long 3’-extensions from their mature forms (12). Moreover, as mentioned above, the use of the conditionally lethal mRNP biogenesis mutants led to the selective production of only aberrant unprocessed precursor mRNAs at the non-permissive condition of 37°C (53–55,57,58,60–63) thereby allowed us to evaluate the consequence of mutation on NRD1 gene on the stability and steady-state levels of these aberrant unprocessed messages. Thus, only the faulty precursor forms of each of the model mRNAs, which specifically accumulate in these mRNP-defective strain backgrounds under the non-permissive condition of 37°C, were stabilized dramatically by the *nrd1-1, nrd1-2* and *nrd1*Δ^CID^ mutants (Figs 1 and 2). This observation thus clearly suggests that Nrd1p facilitates the rapid degradation of these aberrant/precursor forms of these model mRNAs that are generated at the non-permissive temperature of 37°C. Moreover, this view is further strengthened when no significant alteration in the steady-state levels of the test messages in these isogenic yeast strain series was observed at a permissive temperature of 25°C (Figs 1C-D and 2A-D). Strikingly, as noted before (12), while all of these transcripts displayed a similar enhancement in their steady-state levels when exosome component Rrp6p was deleted (in *hpr1*-Δ *rrp6*-Δ and *prp2-1 rrp6*-Δ isogenic strains) and they did not exhibit any stabilization when CTEXT component Cbc1p was deleted (in *hpr1*-Δ *cbc1*-Δ and *prp2-1 cbc1*-Δ isogenic background) (Figs. 1C-D and 2A-B). Strikingly, however, depletion of CTEXT component Cbc1p in the *rna14-1* strain background led to a significant stabilization of two aberrantly long 3’-extended *LYS2* and *CYC1* mRNAs (Fig. 2C-D) (12). Notably, the yeast strains with a defective exosome (*rrp6*-Δ) and CTEXT (*cbc1*-Δ) components were used as controls to assess/compare the efficiency of the Nrd1p-dependent stabilizations of these model aberrant mRNAs.

**Figure 2.**
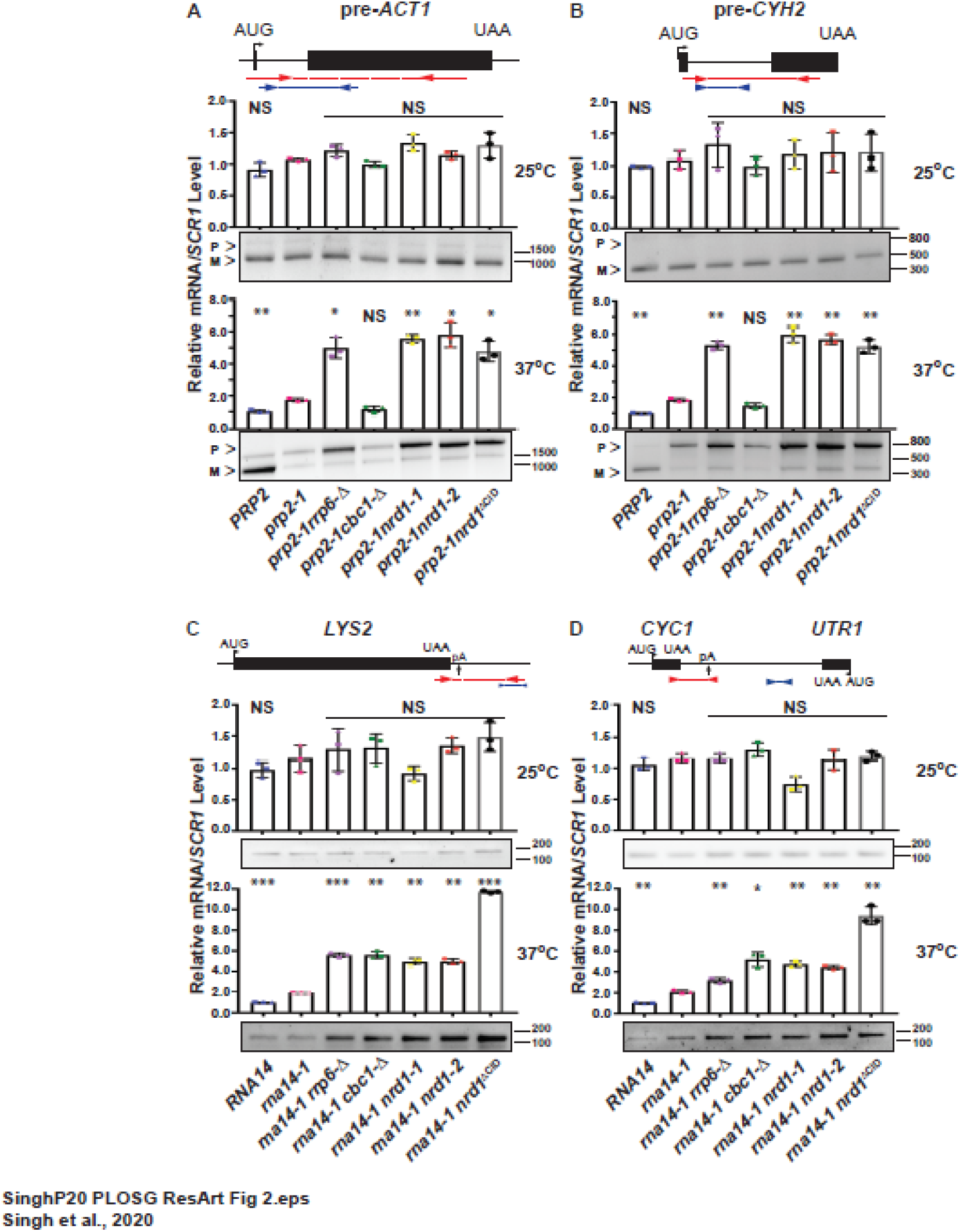
Nrd1p facilitates in the degradation of aberrant mRNPs generated in splice-defective (*prp2-1*) and 3’-end processing-defective (*rna14-1*) yeast strains. Relative abundance of two intron-containing precursor pre-*ACT1* (A) and *pre-CYH2* (B) and two 3’-end processing-defective *LYS2* (C) and *CYC1* (D) mRNAs at permissive (25°C) and non-permissive (37°C) temperature in the *PRP2^+^, prp2-1* and *RNA14^+^, rna14-1* and their indicated isogenic derivative strains as determined by qRT-PCR using a primer set encompassing the amplicon indicated in blue on top of each panel. The normalized value of each pre-mRNA signal from *PRP2*^+^ samples and 3’-end extended transcripts in *RNA14*^+^ strains were set to 1. P-value was estimated with respect to the steady-state levels of the transcript in the *prp2-1* and *rna14-1* strains at each temperature. The panels below each of the graph depicts a representative gel showing the relative levels of the precursor as well as mature *pre-ACT1* and *pre-CYH2* mRNA in A and B and 3’-extended read through transcripts of *LYS2* and *CYC1* mRNAs in C and D as determined by the end-point PCR carried out using primer sets encompassing the amplicons indicated in red on top of each graph. Positions of the relevant size standards are indicated at the right side of each gel panel. **Note that the position of the primer sets used were chosen to specifically distinguish the pre-mRNAs and the read through transcripts from their mature forms and thus enabled us to specifically and precisely estimate only these precursor forms targeted by Nrd1p**. Three independent samples of random-primed cDNA (from biological replicates, N=3) were prepared from indicated strains, pre-grown at 25°C followed by a 3-h shift of the yeast strains at 37°C before subjected to either qRT-PCR or end-point PCR analysis. Transcript copy numbers/50 ng cDNA for *prp2-1* strains and 3 ng for *rna14-1* strains-were normalized to *SCR1* RNA signals obtained from respective samples and are shown as means ± SE. P values were estimated from Student’s two-tailed t-test for a given pair of test strains for each message and is presented with the following symbols, * <0.05, **<0.005 and ***<0.001, NS, not significant.

To further corroborate the enhancement of the steady-state levels of these transcripts in various *nrd1* mutant strains, the decay rates of these faulty messages were determined in wild-type and different *nrd1* strain backgrounds. The decay rates of the selected model mRNAs were determined, post 2 to 3 hours of shifting the growing yeast culture to the non-permissive condition of 37°C followed by the block of RNAPII transcription by 1, 10-phenanthroline and subsequent determination of their steady-state levels at different times by qRT-PCR using the same sets of primer used in the previous experiment. As shown in Figure 3, decay rates of all of the aberrant forms of transcription elongation-defective *ACT1* and *CYC1* mRNAs (Fig. 3A-B), precursor forms of the splice defective pre-*CYH2* mRNA (Fig. 3C) and the faulty read-through transcripts of *LYS2* and *CYC1* mRNAs (Fig. 3D-E) were significantly diminished in *nrd1-1, nrd1-2* and *nrd1*Δ^CID^ strains in comparison to that of the corresponding wild type strains with the concomitant increase in their half-life values (Table 2). Collective data is thus consistent with the conclusion that transcription-elongation/assembly-defective, splice-defective, and aberrantly long read-through transcripts with 3’-extensions are subject to nuclear degradation that is dependent on Nrd1p.

**Figure 3.**
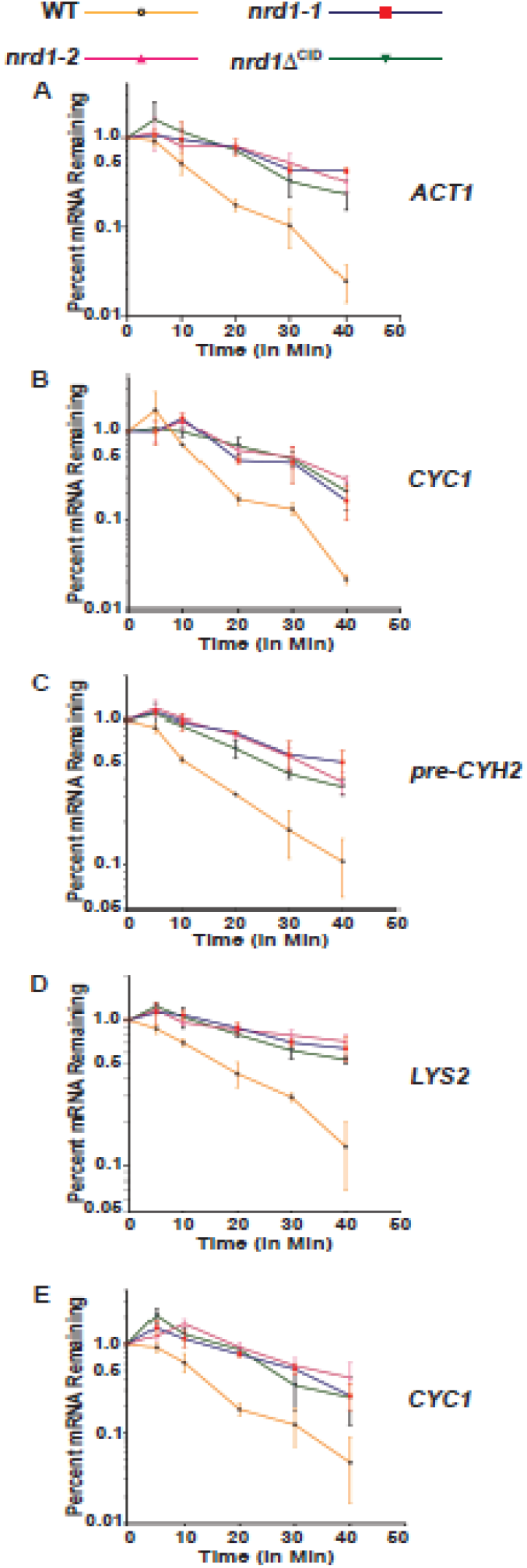
Decay kinetics of representative transcription-elongation-defective *ACT1* and *CYC1* (A-B), splice-defective *pre-CYH2* (C), 3’-end processing-defective *LYS2* and *CYC1* (D-E) mRNAs at 37°C in the wild type (orange line) and yeast strains harboring *nrd1-1* (blue line), *nrd1-2* (pink line), and *nrd1*Δ^CID^ (green line) mutations. Decay rates were determined by qRT-PCR analysis from three independent samples of 2-3 ng cDNA (50 ng for pre-*ACT1* and pre-*CYH2* mRNAs) (biological replicates, N=3) extracted from yeast strains harvested at various times following the treatment of the cells with transcription-inhibitor 100 μg/mL 1, 10-phenanthroline. qRT-PCR signals were normalized to *SCR1* RNA signals obtained from the same cDNA samples, and the normalized signals (mean values ± SD from three biological replicates, N = 3) from each time point were presented as the fraction of remaining RNA (with respect to normalized signals at 0 minute, which was set to 1) as a function of time of incubation in the presence of 1,10-phenanthroline.

**Table 2:** Half-life values of various aberrant mRNAs in different yeast strains carrying various mutant alleles in *NRD1* gene in the backgrounds of transcription-elongation-defect (*hpr1*-Δ), splicing-defect (*prp2-1*) and 3’-end formation (*rna14-1*) defect.

| Yeast Strain defective in | Target aberrant message | Half-life (in minutes) of specific target messages in various *nrd1* mutant strains |  |  |  |
| --- | --- | --- | --- | --- | --- |
|  |  | Wild Type | *nrd1-1* | *nrd1-2* | *nrd1Δ*<sup>CID</sup> |
| **Transcription-Elongation**<br>(*hpr1-Δ*) | *ACT1* | 10.37 | 20.6 | 20.15 | 29.19 |
|  | *CYC1* | 10.22 | 29.94 | 19.77 | 29.96, |
| **Splicing**<br>(*prp2-1*) | pre-*CYH2* | 9.925 | 20.15 | 20.09 | 19.85 |
| **3'-end Processing**<br>(*rna14-1*) | *LYS2* | 10.24 | 20.06 | 19.75 | 19.86 |
|  | *CYC1* | 10.37 | 20.06 | 20.15 | 29.19 |
| **None** | *NCW2* | 10.11 | 19.82 | 19.87 | 9.952 |

A variety of nucleus-retained export inefficient messages are generated during the terminal phase of the mRNP biogenesis in the baker’s yeast which is classified into three different categories, such as (i) those that are generated due to a *cis*-acting mutation in their transcript body, as exemplified by *lys2-187* mRNA (15), (ii) global poly(A)^+^ messages retained in the nucleus of temperature-sensitive *nup116*-Δ mutant yeast strain (14) at the restrictive condition of 37°C due to complete block of nuclear export (65), and (iii) naturally occurring nucleus-retained export-inefficient non-aberrant messages (dubbed special messages) (25) (Table 1). Our previous work established that all of these messages undergo active nuclear degradation by the exosome and its cofactor CTEXT without any participation of TRAMP (12). An assessment of the functional role of Nrd1p in the decay of export-incompetent messages collectively revealed that all the model aberrant export-defective mRNAs were stabilized dramatically in *nrd1-1* and *nrd1-2* strain backgrounds (Fig 4–5). The destabilization of *lys2-187* mRNA in *lys2-187* isogenic strains at 25°C (Fig 4B), nucleus-arrested *ACT1* and *CYC1* mRNA in *nup116*-Δ isogenic strains at 37°C (Fig 4E and G) and three export-incompetent special mRNAs *NCW2, SKS1* and *IMP3* in normal isogenic strains at 25°C (Fig 5B-D) were all rescued in the corresponding isogenic *rrp6-Δ, cbc1-Δ, nrd1-1*, and *nrd1-2* mutant yeast strains. As expected, the negative controls for this experiment, normal export-efficient *CYC1* mRNA in *lys2-187* isogenic series (Fig. 4A), *ACT1* and *CYC1* mRNA in *nup116*-Δ isogenic strains at 25°C (Fig 4D and F), and *CYC1* in isogenic strain background with no mRNA biogenesis defect (Fig. 5A) did not display in any significant destabilization and concomitant stabilization by the *nrd1* mutations. Surprisingly and remarkably, none of the export-defective test messages displayed any steady-state enhancement at all in *nrd1*Δ^CID^ mutant isogenic strain (Fig 4–5). Consistent with the pattern of steady-state enhancement of the mutant *lys2-187* message in the *nrd1-1* and *nrd1-2* mutant strains, *lys2-187* strain also displayed a compromised growth in the SC-*lys* medium as compared to normal *LYS*^+^ strain, which was rescued in all of the *lys2-187 nrd1-1, lys2-187 nrd1-2, lys2-187 rrp6*-Δ, *lys2-187 cbc1*-Δ yeast strains but not in a *lys2-187 nrd1ΔCID* strain (Fig 4C). Furthermore, the decay rate of *NCW2* message was found to be significantly (2-3 folds) diminished in *nrd1-1, nrd1-2* strains with concomitant increase in half-life values when compared to its decay rates in the wild-type and *nrdl*Δ^CID^ strains (Table 2, Fig 5E) thereby nicely correlated to the steady-state enhancement profile of this special message in these strains. As discussed later in the results and discussion, the peculiar behavior of the *NCW2* mRNA and other export defective messages in general was correlated to the non-essentiality of the requirement of the CID domain of the Nrd1p, since the nrd1Δ^CID^p is equally capable of supporting the rapid degradation of the *NCW2* and other export inefficient messages (Fig. 4B, 4E, 4G, 5B-E). Collectively, thus, these data suggest that Nrd1p participates in the nuclear decay of all classes of export-defective transcripts. Interestingly, while the *nrd1-1, nrd1-2* mutations enhanced the steady-state levels of all of the export-inefficient transcripts, the *nrd1*Δ^CID^ mutation did not enhance their abundance at all, thereby supporting the notion that *nrd1*Δ^CID^ protein is as efficient in supporting the degradation of the export defective messages as the wild type Nrd1p.

**Figure 4.**
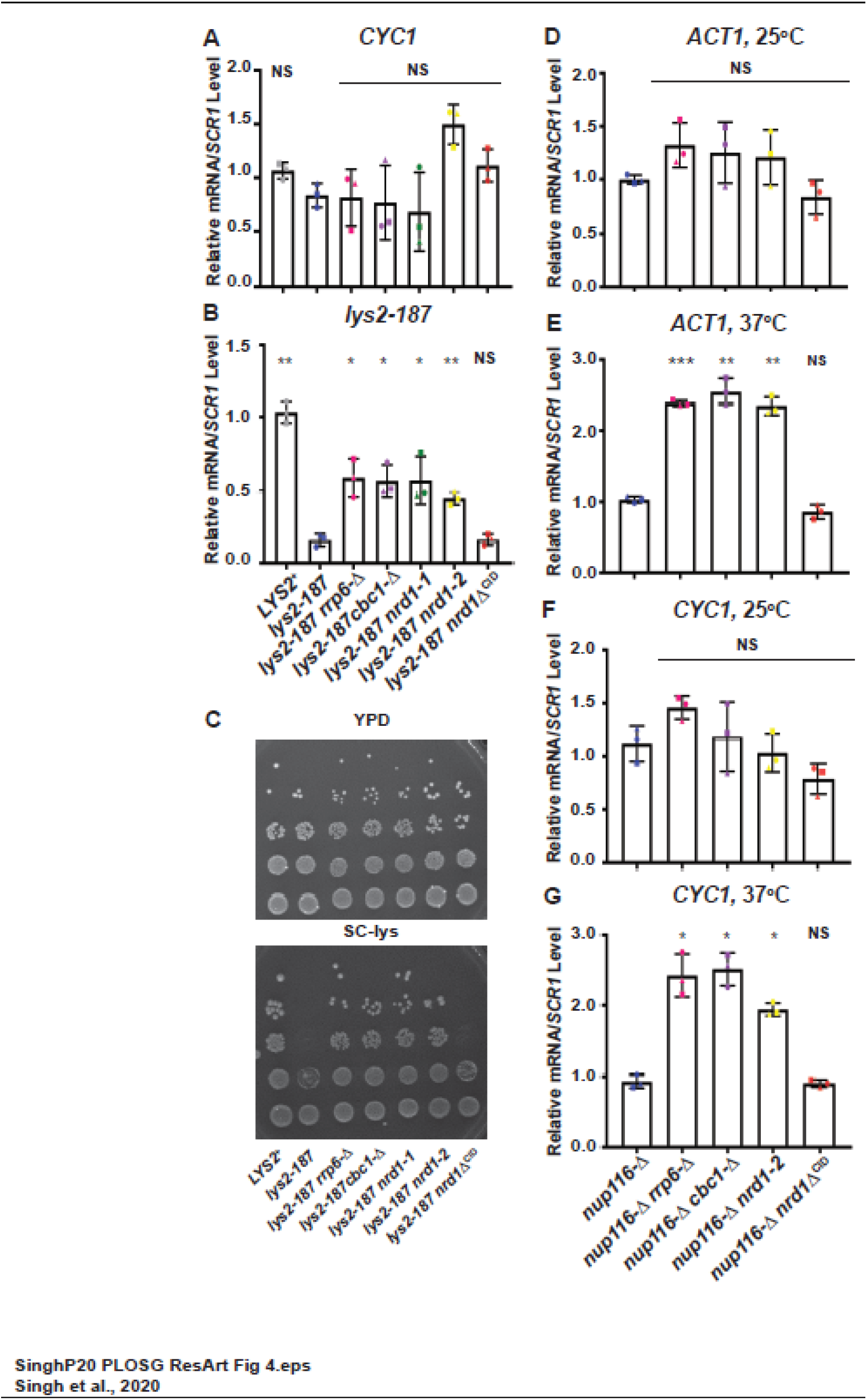
Rapid nuclear decay of export-defective messages is rescued by *nrd1-1* and *nrd1-2* mutations, but not by *nrd1*Δ^CID^ mutation. (A-B) Relative steady-state levels of export efficient normal *CYC1* (negative control) (A) and normal *LYS2* and export-defective *lys2-187* (B) mRNAs at 25°C in the indicated isogenic yeast strains. The normalized value of the *CYC1* and *LYS2* transcripts from the *LYS2*^+^ sample was set to 1. P-value was estimated with respect to the steady-state levels of the respective transcripts in the *lys2-187* yeast strain (A-B). C. Growth profile of *LYS2*^+^ and the isogenic *lys2-187* mutant strains on SC-Lys medium at 25°C. A series of progressively diluted (1/10^th^) suspension from the stock (concentration 10^5^/cells per ml) of each strain were made, followed by spotting (10 μl) on lysine omission medium (SC-Lys). The plates were incubated at 25°C for 3-4 days before the capturing the images. (D-G) Relative steady-state levels of nucleus-retained *ACT1* (D-E) and *CYC1* (F-G) at permissive (25°C) and non-permissive (37°C) in the *nup116*-Δ mutant and its various indicated isogenic strains. The normalized value of these transcripts from the *nup116*-Δ sample was set to 1. Transcript copy numbers/3 ng cDNA for *lys2-187* strains, copy numbers/2 ng cDNA of other strain backgrounds were normalized to *SCR1* RNA levels in respective samples and are shown as means ± SE. P values estimated from Student’s two-tailed t-test for a given pair of test strains for each message, is presented with the following symbols, * <0.05, **<0.005 and ***<0.001, NS, not significant.

**Figure 5.**
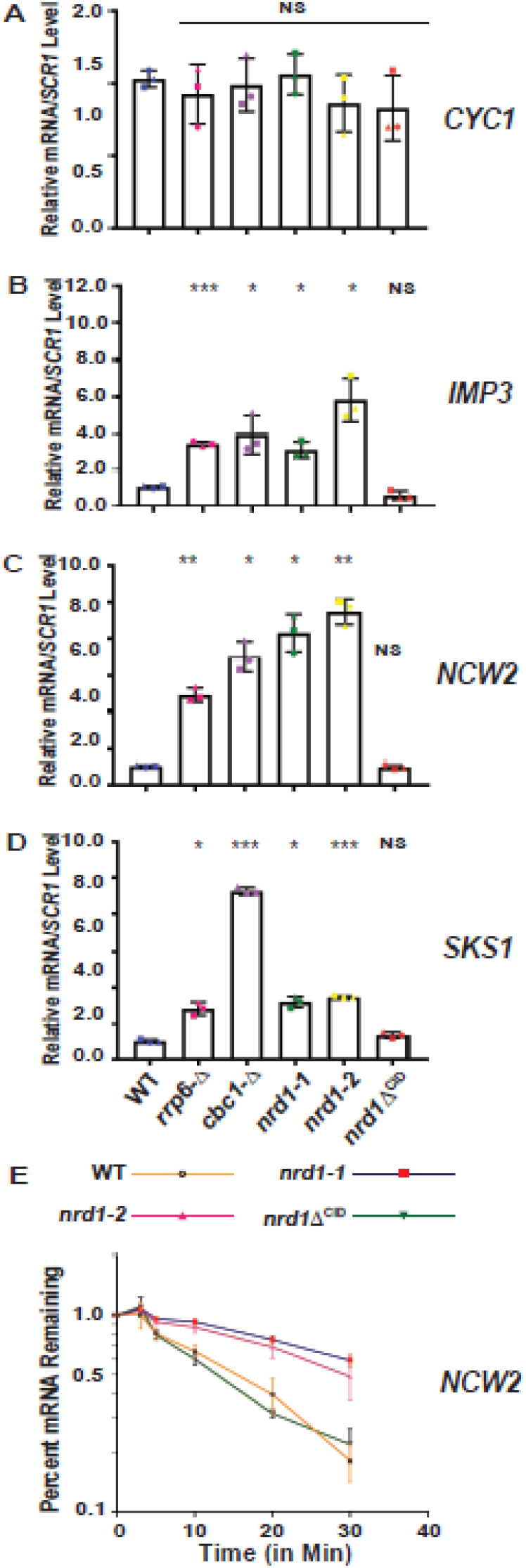
*nrd1-1* and *nrd1-2*, but not *nrd1*Δ^CID^ mutation rescue the rapid nuclear degradation of export-inefficient nucleus-retained special messages. (A-E) Relative steady-state levels of one export efficient mRNA *CYC1* (A) and three export-inefficient special mRNAs, *IMP3* (B), *NCW2* (C) and *SKS1* (D) at 25°C in a wild type, *rrp6*-Δ, *cbc1*-Δ, *nrd1-1, nrd1-2*, and *nrd1*Δ*CID* strains. The normalized value of each specific mRNAs from the wild-type sample was set to 1. Three independent samples of random-primed cDNA (biological replicates, N = 3) samples were prepared from the indicated isogenic strains grown at 25°C before subjecting them to real-time qPCR analysis using primer sets specific for each mature mRNA. Transcript copy numbers/3 ng cDNA were normalized to *SCR1* RNA levels in respective samples and are shown as means ± SE. P values estimated from Student’s two-tailed t-test for a given pair of test strains for each message, is presented with the following symbols, * <0.05, **<0.005 and ***<0.001, NS, not significant. (F) Decay kinetics of *NCW2* mRNA at 25°C in the wild type (orange line) and yeast strains harboring *nrd1-1* (blue line), *nrd1-2* (pink line), and *nrd1*Δ^CID^ (green line) mutations. Decay rate was estimated after addition of transcription inhibitor 1,10-phenanthroline as described before.

Next, we established genetic epistasis between the core exosome/CTEXT components, Rrp6p, Cbc1p, and NNS component, Nrd1p, while evaluating if the core-exosome, CTEXT and NNS complex collectively act as a part of the same pathway. Findings from this experiment revealed that steady-state levels of the two special messages, *NCW2* and *SKS1* observed in the single mutant *rrp6*-Δ, *cbc1*-Δ and *nrd1-2* strains, did not further enhance in the double mutant *rrp6*-Δ *nrd1-2, cbc1*-Δ *nrd1-2* strains (Fig 6B-C). Finally, while evaluating if the entire NNS complex (Nrd1p-Nab3p-Sen1p) collectively participates in the degradation of aberrant export-defective transcripts, we also observed that the steady-state levels of the special mRNAs, *IMP3, NCW2*, and *SKS1* mRNAs were dramatically stabilized in the yeast strains carrying *nab3-10* (four to eight folds) (Fig 6E-G) and *sen1-1* (two to five folds) (Fig. 6I-K) mutations as compared to wild type strain. The normal *CYC1* mRNA, in contrast, neither displayed any destabilization nor stabilization in any of these strains (Fig 6A, D, H). This data is thus consistent with the conclusion that Nrd1p-Nab3p-Sen1p together plays a crucial functional role in the decay of export defective special messages. Collectively, therefore, the findings from the experiments presented in this section, very strongly suggest that Nrd1p-Nab3p-Sen1p complex constitutes an integral component of the nuclear degradation machinery that targets all types of aberrant nuclear mRNAs (Table 3).

**Figure 6.**
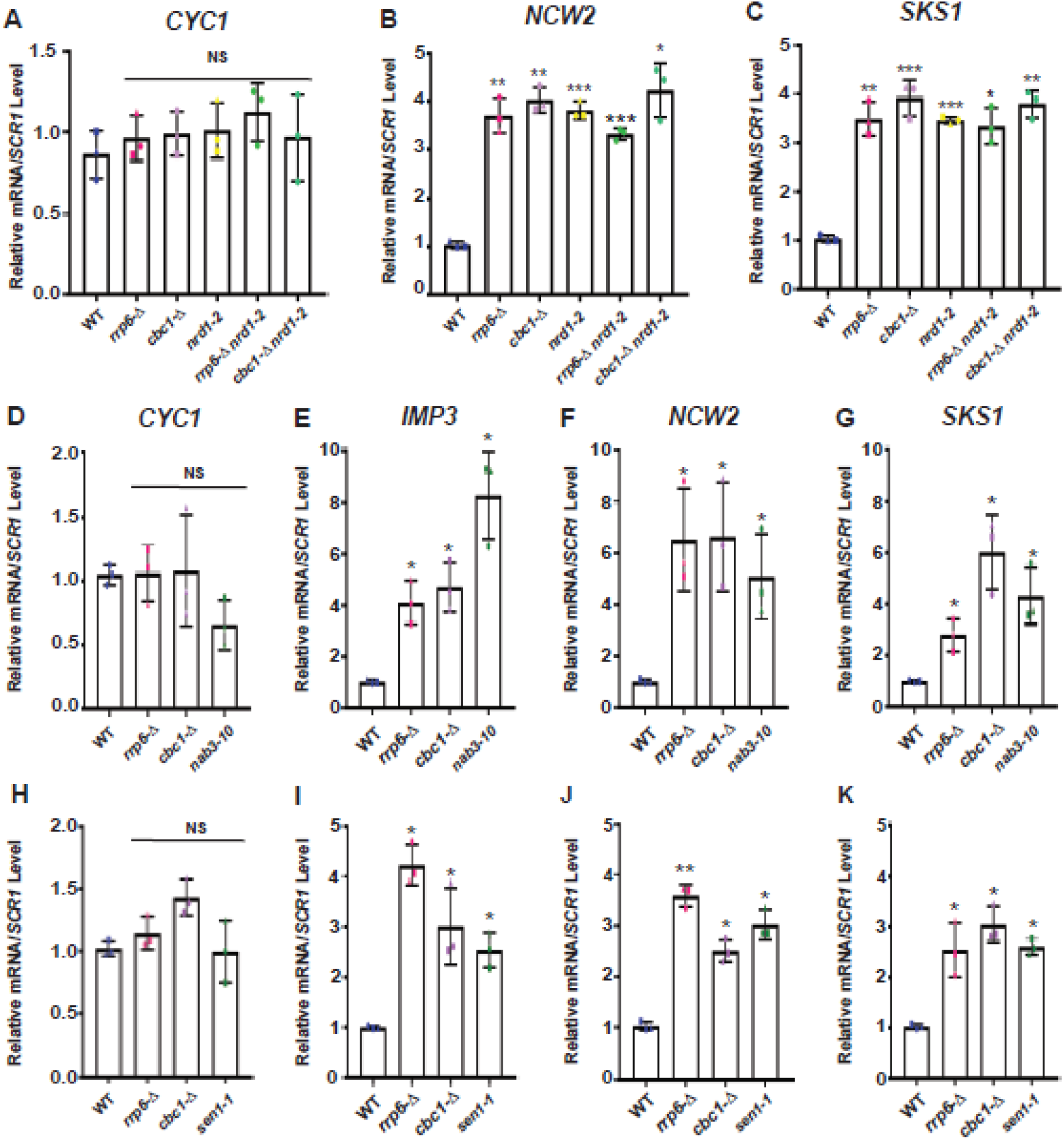
All the components of NNS complex act together with the core nuclear exosome, and CTEXT complex to facilitate the decay of the export defective special mRNAs. (A) Relative steady-state levels of export efficient *CYC1* mRNA, and (B-C) two export-inefficient special messages, *NCW2* (B) and *SKS1* (C) mRNAs at 25°C in an isogenic wild type, *rrp6*-Δ, *cbc1*-Δ, *nrd1-2, rrp6*-Δ *nrd1-2*, and *cbc1*-Δ *nrd1-2* strains. The normalized value of each specific mRNAs from the wild-type sample was set to 1. (D-K) Relative steady-state levels of normal export-efficient *CYC1* (D and H) and three export-inefficient special mRNAs, *IMP3* (E and I), *NCW2* (F and J) and *SKS1* (G and K) at 25°C in the indicated *nab3* (D-G), and *sen1* (H-K) isogenic yeast strain series. Normalized values of the transcript in the wild-type samples were set to 1. The RNA extraction and steady-state level of the specific mRNAs were determined as described in the legend of Figure 1. P values were estimated from Student’s two-tailed t-test for a given pair of test strains for each message and is presented with the following symbols, * <0.05, **<0.005 and ***<0.001, NS, not significant.

**Table 3:**
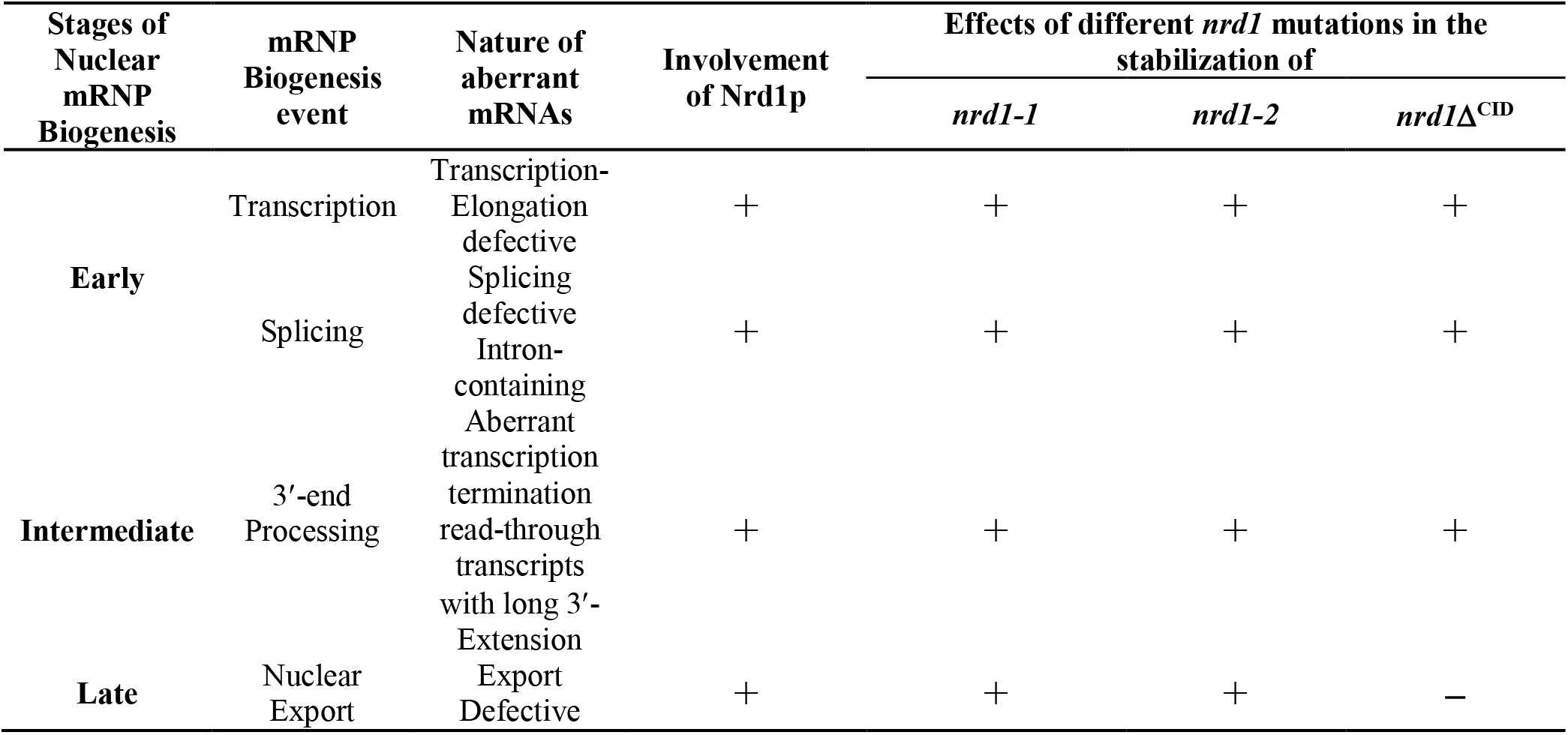
Specificity of Nrd1p in the decay of various aberrant nuclear mRNA targets examined in this study and the effect of various *nrd1* mutants in their stabilization.

Strikingly, the *nrd1*Δ^CID^ mutation, unlike the *nrd1-1* and *nrd1-2* mutations, displayed a differential specificity towards the aberrant nuclear transcripts. While a dramatic effect of the *nrd1*Δ^CID^ mutant was observed in stabilizing the transcription/assembly-, splice-, and 3’-processing-defective mRNAs (Figs 1–3), no stabilizing influence of this mutant was noted in case of export defective messages (Figs 4–5) (Table 3). This observation suggests that although *nrd1*Δ^CID^p is incapable of supporting the decay of the aberrant messages produced in the early and intermediate stages of mRNP biogenesis, it is able to promote the rapid decay of the export-incompetent messages. Since the interaction of Nrd1p-RNAPII is abolished in the *nrd1*Δ^CID^ strain (18,20,30,33,66), our data strongly indicate that the efficient degradation of aberrant mRNAs generated during transcript-elongation, splicing and 3’-end formation (early and intermediate stage of mRNP biogenesis) requires Nrd1p complex to be recruited co-transcriptionally onto these transcripts via RNAPII-CTD. For the rapid degradation of export-defective messages, in contrast, the CID domain appears dispensable, leading to the conclusion that nrd1Δ^CID^p still supports the nuclear decay of the aberrant export-incompetent messages in an RNAPII-CTD independent manner. This conclusion, therefore, prompted us to look into the molecular insight into the co-transcriptional recruitment profile of the Nrd1p on various kinds of faulty messages to see if its co-transcriptional recruitment onto a specific faulty message is vital for its nuclear degradation.

### RNAPII-dependent co-transcriptional recruitment of Nrd1p is vital for the exosome/TRAMP-dependent decay of the aberrant mRNAs derived during the early and intermediate stages of mRNA biogenesis

We begin this section with a naïve question; if Nrd1p recruitment on a faulty message is significantly higher in comparison to its normal functional counterpart. Consequently, the Nrd1p occupancy profiles of the *ACT1* and *CYC1* messages in *hpr1*-Δ mutant strains at 37°C as determined by chromatin immunoprecipitation (ChIP) were found to be dramatically higher than those estimated at 25°C (Fig 7A-B). Similarly, the Nrd1p occupancy of the export-incompetent *NCW2* mRNA was significantly higher in comparison to typical export-efficient message *CYC1* (Fig. 7C). This data supports the conclusion that perhaps Nrd1p (and NNS complex) is selectively recruited on to faulty and other target RNAs (such as special messages) of the nuclear exosome in a co-transcriptional manner. Next, we determined the occupancy profiles of C-terminally truncated Nrd1p in yeast strains carrying the *nrd1-1* and *nrd1-2* alleles, which were dramatically low on the *ACT1* and *CYC1* in *hpr1*-Δ isogenic yeast strain background at 37°C as well as on the *NCW2* and *SKS1* messages in wild-type yeast strain background (Supplementary Fig. S1). This low level of recruitment could be the result of the inability of the nrd1-1p and nrd1-2p mutant proteins lacking RNA recognition motif to bind to these RNAs (18,33). Alternatively, the expression of these mutant proteins in the *nrd1-1* and *nrd1-2* yeast strains were possibly very low(20), leading to their weak association with these mRNAs. Thus, the low co-transcriptional recruitment obtained for the truncated nrd1-1p and nrd1-2p prompted us to restrict ourselves in evaluating the Nrd1p occupancy profile in *NRD1*^+^ and *nrd1*Δ^CID^ mutant yeast strains only for all the subsequent co-transcriptional recruitment experiments. Nevertheless, our data (i) indicates that Nrd1p recruitment onto aberrant messages perhaps aids in the initial recognition of these targets and (ii) hints at the possibility that during the transcription elongation the primary distinction between the functional vs. aberrant messages is probably accomplished by the presence of Nrd1p on these faulty transcripts.

**Figure 7.**
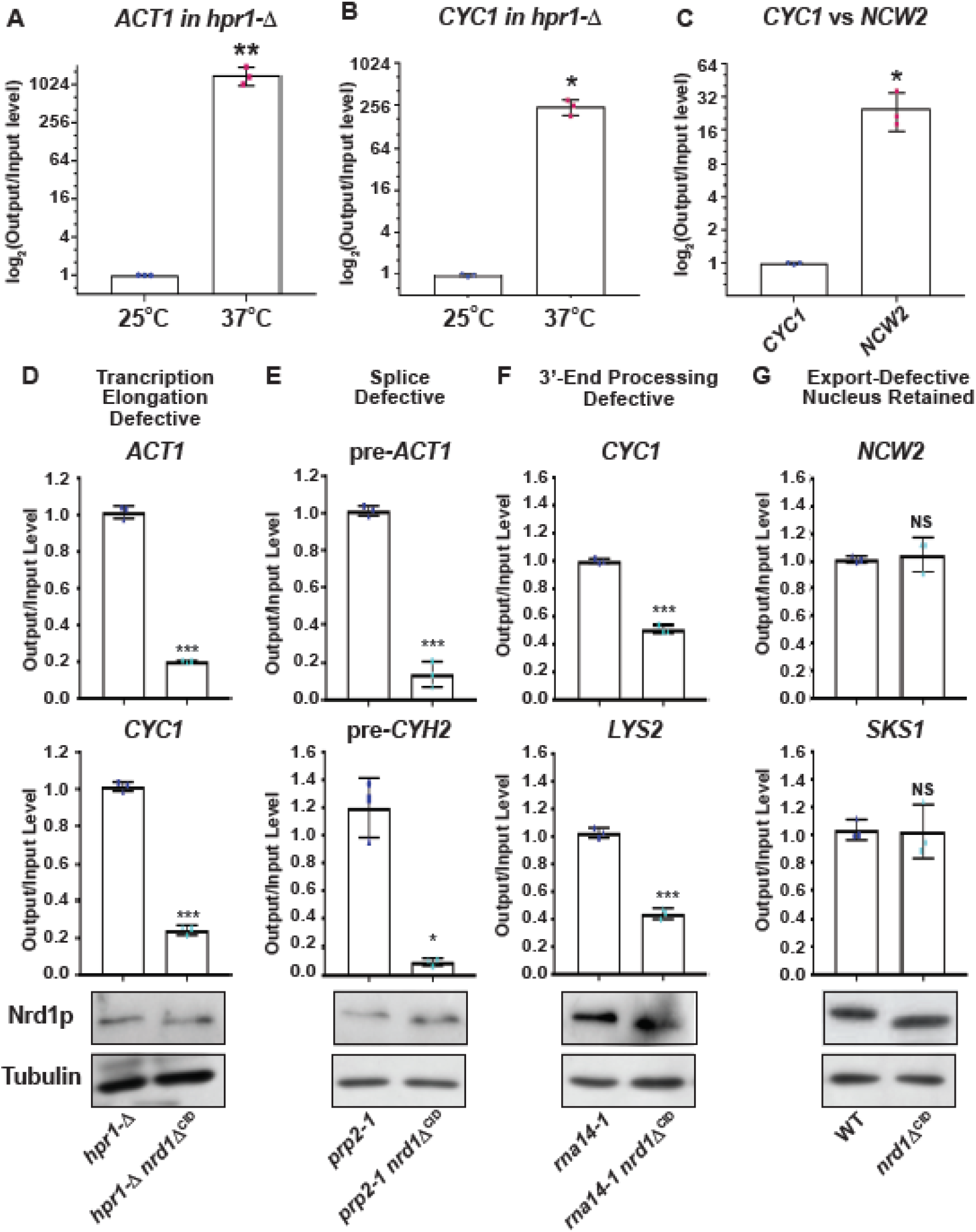
The co-transcriptional recruitment of Nrd1p lacking the *CID* domain is drastically impaired on transcription assembly-defective, splice-defective, and 3’-extended aberrant messages in comparison to full-length Nrd1p but remains unaltered on export-defective special messages. (A-C) Nrd1p occupancy profiles of the transcription-assembly-defective *ACT1* (A) and *CYC1* (B) in an *hpr1*-Δ strain at 25°C and 37°C, and a normal export efficient, *CYC1* and special *NCW2* mRNAs (C) in a wild-type yeast strain at 25°C. (D-G) Occupancy profile of full-length Nrd1p and Nrd1p lacking its *CID* domain of the (D) transcription-assembly-defective *ACT1* and *CYC1* mRNAs at 37°C in isogenic *hpr1*-Δ and *hpr1*-Δ *nrd1*Δ^CID^ strains, (E) splice-defective intron-containing pre-*ACT1* and pre-*CYH2* at 37°C in isogeneic *prp2-1* and *prp2-1 nrd1*Δ^CID^ strains (F) 3’-extended read-through *CYC1* and *LYS2* transcripts at 37°C in *rna14-1* and *rna14-1 nrd1ΔCID* strains and (G) nucleus-arrested special messages *NCW2* and *SKS1* at 25°C in wild-type (*NRD1*^+^) and *nrd1ΔCID* strains. Panels at the bottom of each graph depicts the expression levels of Nrd1p (both full-length and CID-deleted version) and tubulin as determined by western blot using anti-Nrd1p and anti-tubulin antibodies in each of the isogenic yeast strain series harboring either an *NRD1*^+^ or an *nrd1*Δ^CID^ allele. Each lane contained an equal amount of quantified protein extracts from the indicated strains. A 70 kDa band in the extract from *NRD1+* strain and 55 kDa band in the extract from *nrd1*Δ^CID^ was detected with no non-specific band. For ChIP assays, fragmented and cross-liked chromatin preparation was conducted from indicated strains at a specific temperature from three independent biological replicates (N=3) followed by the immunoprecipitation of the chromatin samples using specific Nrd1p antibody. Immunoprecipitated DNA was recovered as mentioned in materials and methods before qPCR analyses using the primer sets specific for the middle region of ORF of each target mRNA. Mean normalized qPCR signals ± SD from the immunoprecipitated (output) DNA to the mean qPCR signal obtained from the total chromatin (input) DNA of each sample obtained from three experiments are presented. P-values estimated from Student’s two-tailed t-tests for a given pair of test strains for each message, is presented with the following symbols, * <0.05, **<0.005 and ***<0.001, NS, not significant.

Consequently, we determined the Nrd1p recruitment profile onto the two representative model mRNAs from every class of aberrant messages in isogenic yeast strains carrying either an *NRD1*^+^ or *nrd1*Δ^CID^ allele. We generated the Nrd1p occupancy profile of each mRNA using the primer sets located at the 5’-end, middle, and at the 3’-termini, which were essentially very similar. For clarity, the occupancy profile obtained with the primer sets corresponding to the central segment of each of the target mRNAs is presented in Fig 7D-G. Our findings revealed that, the representative transcription-elongation defective mRNAs in *hpr1*-Δ *NRD1*^+^ strain, splice defective mRNAs in *prp2-1 NRD1*^+^ strain and 3’-end processing-defective transcripts in *rna14-1 NRD1*^+^ strain at the restrictive temperature of 37°C display a significantly high occupancy of Nrd1p in comparison to that estimated in corresponding *nrd1*Δ^CID^ isogenic strains (Fig 7D-F). Furthermore, low recruitment of Nrd1p observed in these *nrd1*Δ^CID^ strains was not associated with its low expression level as demonstrated by the similar and comparable Nrd1p expression levels in both *NRD1*^+^ and *nrd1*Δ^CID^ yeast strains as shown in the bottom panels of Fig 7D-F. This observation thus clearly indicated that Nrd1p occupancy onto the test mRNAs in an *nrd1*Δ^CID^ yeast strain is significantly lower because of its lower co-transcriptional recruitment, owing to the lack of association of N-terminally truncated nrd1Δ^CID^p protein with the RNAPII-CTD. This conclusion thereby justifies the view that Nrd1p recruitment on these aberrant messages derived in the early and intermediate phase of mRNP biogenesis is dependent on the RNAPII-CTD, which is vital for marking these messages as faulty.

### Nuclear degradation of export-defective aberrant mRNAs generated in the late phase of mRNP biogenesis requires Pcf11p-dependent recruitment of Nrd1p that precedes the recruitment of the Rrp6p/nuclear exosome

Finally, we determined that the Nrd1p occupancy on the two representative export-defective special messages, *NCW2* and *SKS1* mRNAs in the wild type and *nrd1*Δ^CID^ mutant yeast strains at 25°C was very similar (Fig 7G). This observation strongly supported the view that Nrd1p recruitment on export-defective messages (i) took place normally in an *nrd1*Δ^CID^ strain and supported their rapid decay, and (ii) does not require its CID domain, and hence it is independent of the RNAPII-dependent recruitment. Collectively, thus, we establish that Nrd1p (and NNS complex) participates in the nuclear degradation of all kinds of aberrant mRNA targets derived at various phases of mRNP biogenesis and its recruitment onto these aberrant messages is vital for their nuclear decay. Interestingly, however, the co-transcriptional recruitment of Nrd1p in the early/intermediate phase is strongly dictated by its interaction with RNAPII (via the interaction of the RNAPII CTD-Nrd1pCID domain). In contrast, its recruitment onto the defective messages generated at the terminal phase of mRNP biogenesis is independent of RNAPII. To find out the cellular recruiter of Nrd1p onto the export-inefficient messages, we scrutinized the literature in search of an ideal factor, which should be characterized by its ability (i) to support only the decay of export-defective messages but not the decay of other aberrant classes, (ii) to selectively promote the recruitment of Nrd1p on the model-export defective messages only, and (iii) to physically interact with Nrd1p.

A thorough search prompted us to postulate that, Pcf11p, a key component of cleavage and polyadenylation specific (CPSF) complex, which is involved in the transcription termination of poly(A)^+^ RNAs may serve as a putative recruiter. Remarkably, Pcf11p also possesses a CID (CTD-interacting domain) like Nrd1p, and this protein interacts with RNAPII and many nascent mRNAs after being recruited by RNAPII via its CID (67–71). Most interestingly, Nrd1p, and Pcf11p were found to interact physically by affinity capture RNA method (44) and genetically by dosage rescue (72). Our initial effort to test the Pcf11p as the possible recruiter of Nrd1p revealed that a mutant yeast strain harboring a *pcf11-2* allele (67) did not enhance the steady-state levels of two representative transcription-elongation/assembly-defective messages in THO mutant isogenic strain background (Fig 8A). In contrast, the *pcf11-2* allele firmly stabilized the two export-defective special messages, *NCW2* and *SKS1* mRNAs (Fig 8B), thereby strongly supporting our prediction. Consistent with this finding, a comparison of the Nrd1p occupancy profiles on the same group of representative messages displayed that, while recruitment profile on the transcription-elongation assembly defective *ACT1* and *CYC1* messages remained unaltered in *pcf11-2* mutant yeast strain, it is significantly reduced on the *NCW2* and *SKS1* mRNAs in the same *pcf11-2* strain (Fig 8C-D). This finding is thus consistent with the conclusion that Pcf11p plays a vital functional role in the co-transcriptional recruitment of Nrd1p onto the export-defective messages, thereby promoting their nuclear degradation by the Nrd1p/nuclear exosome.

**Figure 8.**
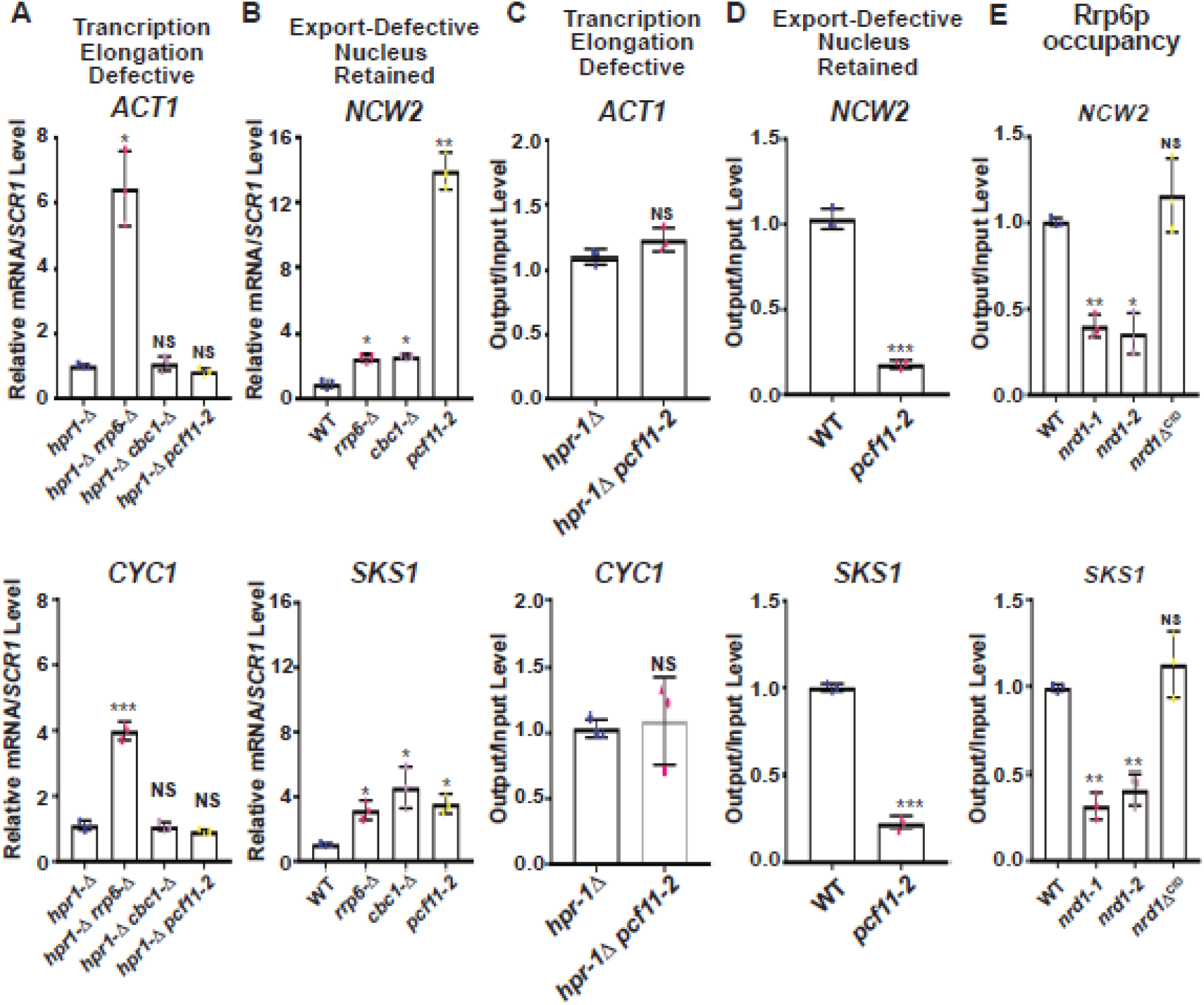
Pcf11p actively contributes to the rapid nuclear decay of export-defective special messages by recruiting the Nrd1p that leads to the further recruitment of Rrp6p onto them. (A-B) Relative steady-state levels of (A) two transcription assembly-defective *ACT1* and *CYC1* mRNAs in the indicated isogenic *hpr1*-Δ, *hpr1*-Δ *rrp6*-Δ, *hpr1*-Δ *cbc1*-Δ and *hpr1*-Δ*pcf11-2* strains and (B) two export-inefficient special mRNAs *NCW2* and *SKS1* in isogenic wild-type (*PCF11*^+^), *rrp6*-Δ, *cbc1*-Δ and *pcf11-2* strains. In both cases, the normalized values of every target mRNA from the wild-type samples was set to 1. Normalized qRT-PCR signals (Mean ± SE) from three independent biological replicates are presented as scatter plots, as described above. (C-D) co-transcriptional recruitment profiles of Nrd1p on (C) transcription assembly-defective *ACT1* and *CYC1* messages at 37°C in isogenic *hpr1*-Δ and *hpr1-Δ pcf11-2* strains, and (D) two export-incompetent special messages in *NCW2* and *SKS1* at 25°C in an isogenic wild-type and *pcf11-2* strains. (E) Occupancy profiles of Rrp6p on the two special mRNAs *NCW2* and *SKS1* at 25°C in the indicated isogenic wild-type (*NRD1*^+^), *nrd1-1, nrd1-2*, and *nrd1*Δ^CID^ yeast strains. All immunoprecipitations using specific Nrd1p or Rrp6p antibody followed by the recovery of the precipitated DNA and qPCR analyses were performed exactly as described in materials and methods and the legend of Figure 6. P-values estimated from Student’s two-tailed t-tests for a given pair of test strains for each message, is presented with the following symbols, * <0.05, **<0.005 and ***<0.001, NS, not significant.

Next, we demonstrated that the Nrd1p recruitment on the aberrant export defective mRNAs leads to the further recruitment of the exosome component Rrp6p onto export-incompetent messages by showing that Rrp6p occupancy profiles on the two special mRNAs, *NCW2* and *SKS1* is very high in both the *NRD1*^+^ and *nrd1*Δ^CID^ mutant strains, whereas, it is significantly low in *nrd1-1* and *nrd1-2* strains (Fig 8E). This data thus strongly supports our conclusion that co-transcriptional recruitment of Nrd1p onto the export-defective aberrant messages (mediated by Pcf11p) is followed by the subsequent recruitment of the nuclear exosome, thereby stimulating their rapid nuclear decay. Thus, collectively, our experimental findings are consistent with the conclusion that the trimeric Nrd1p-Nab3p-Sen1p complex constitutes a vital and universal component of the nuclear decay and surveillance apparatus in *Saccharomyces cerevisiae* and its co-transcriptional recruitment is crucial for the nuclear degradation of the entire spectrum of aberrant mRNAs.

### Several export-incompetent nucleus-retained special mRNAs naturally harbor Nrd1p RNA-binding motif in their coding regions and have been demonstrated to bind Nrd1p under normal conditions

Previous analysis of the Nrd1p PAR-CLIP data revealed that Nrd1p binds to the known UGUAG motif (73–75) and this site is strongly overrepresented in the 3’-end of the non-coding RNA genes but is generally lacking in the mRNAs except for those whose expression are regulated by Nrd1p-Nab3p-Sen1p via transcription termination (22,73–75). Our findings that the export-defective special mRNAs are targeted by Nrd1p-dependent nuclear decay under normal condition in the wild-type yeast strain, prompted to us to verify if any of these mRNAs harbor any putative tetrameric Nrd1p-binding site(s) and actually bind Nrd1p *in vivo*. To address this question, we mined the previously published Nrd1p PAR-CLIP dataset GSE31764 (gene Expression Omnibus (http://www.ncbi.nlm.nih.gov/geo/query/acc.cgi?acc=GSE31764) and analyzed the processed wig and FASTA files associated with this datasets (73). As shown in Fig 9C-D, at least two of the three experimentally tested special mRNAs, *NCW2* and *IMP3* display strong Nrd1p cross-linked sites in their coding region. Moreover, the Nrd1p-motif perfectly overlaps with the sites of T→C transition thereby bolstering that notion that Nrd1p actually binds to its perfect GUAG motif *in vivo* for both of these mRNAs (Fig 9C-D). In contrast, the wig files show no Nrd1p binding sites in the coding regions of two typical mRNAs, *ACT1* and *CYC1* (Fig 9A-B). It is noteworthy to mention here that we did not expect to find strong mRNA binding motifs in the coding region of any of the model representative mRNAs, *ACT1, CYC1*, and *LYS2* etc., which were used in our analysis of steady-state and co-transcriptional recruitment analysis in the background of various mRNP biogenesis mutants as they are not targeted by Nrd1p under permissive condition. We believe that the nature of co-transcriptional recruitment/binding of Nrd1p onto these messages under the conditional temperature of 37°C is different from its binding to the specific Nrd1 target messages (such as *NRD1, PCF11, NCW2* and *IMP3* mRNAs etc.) (See discussion).

**Figure 9:**
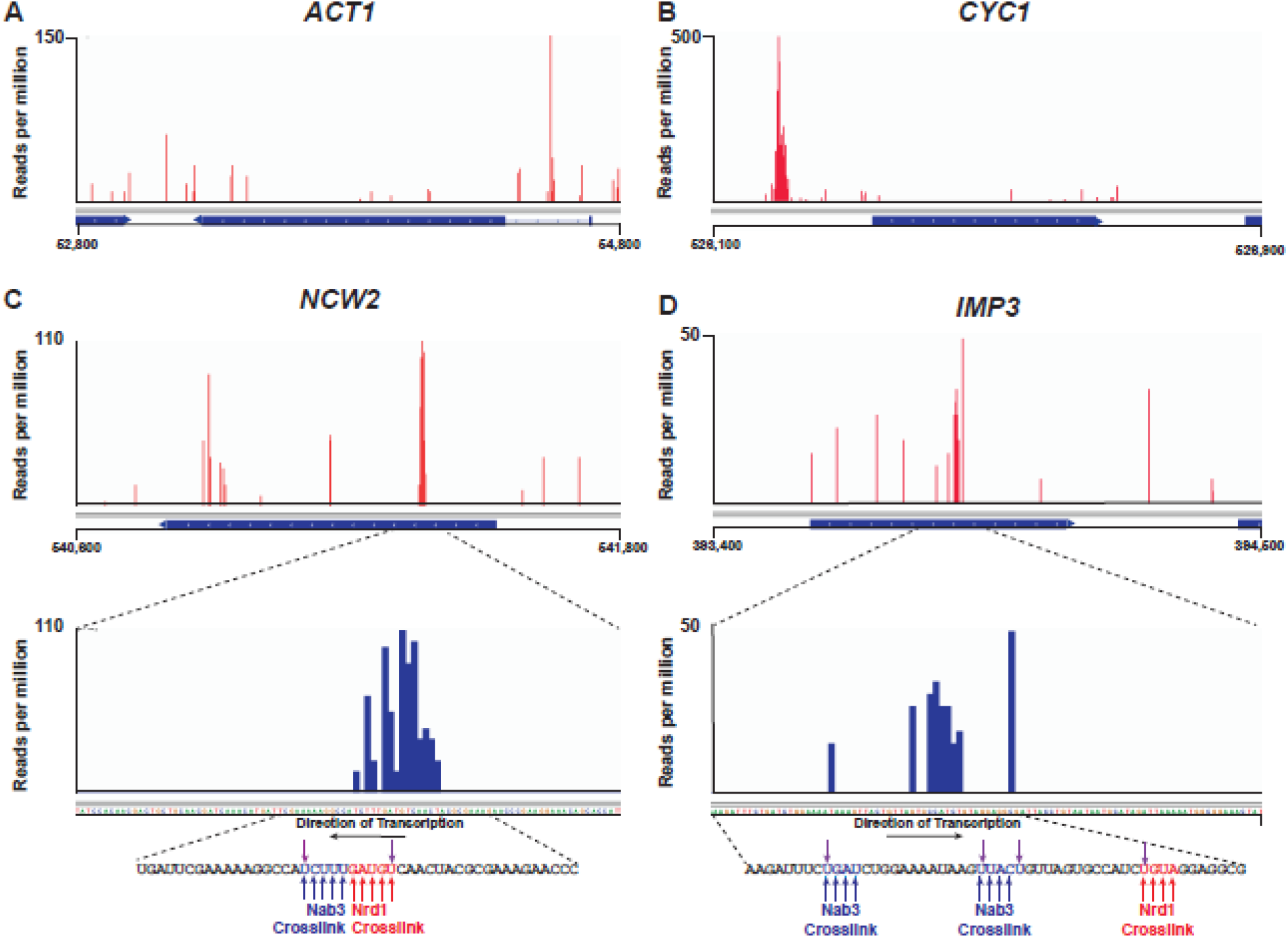
*In vivo* Nrd1p and Nab3p cross linking to *NCW2* and *IMP3* RNAs. Cross-linking and binding profiles of Nrd1p from (A) a 2000 nucleotide genomic region of chromosome VI harboring *ACT1;* (B) 800 nucleotide region of chromosome X harboring *CYC1*, (C) 1200 nucleotide region of chromosome XII harboring *NCW2*, and (D) 1100 nucleotide region of chromosome VIII harboring *IMP3* gene. A higher-resolution map of cross-linked reads to the *NCW2* and *IMP3* coding region are shown at the bottom of the original reads. The number of reads per 10^7^ reads is indicated on the y-axis. Sequences of the major cross-linked regions at the bottom of the figure. The upwards arrows correspond to the Nrd1 (blue) or Nab3 (red) cross-linking motifs and the downward purple arrows indicate the sites most frequently underwent T→C transitions in both Nrd1 and Nab3p data sets.

Thus the experimental evidence presented above collectively affirm that Nrd1p-Nab3p-Sen1p complex constitutes a pivotal component of the nuclear surveillance mechanism in *Saccharomyces cerevisiae* and Nrd1p binding to aberrant messages is vital for their nuclear decay. This conclusion is further supported by the finding that at least for the aberrant export defective messages, the co-transcriptional recruitment of this complex promotes the subsequent recruitment of the nuclear exosome for facilitating their degradation. Thus, the Nrd1p-Nab3p-Sen1p complex appears to serve as an exosome-specificity factor that co-transcriptionally recognizes the aberrant mRNAs very early in the transcription and mRNP biogenesis.

## Discussion

Although our previous work demonstrated a differential distribution of duty among the two exosomal cofactors, TRAMP and CTEXT, and their stage-specific recruitment on the diverse faulty messages (12), the mechanism of stage-specific recognition of distinct aberrant message is largely unknown. Moreover, although the existence of an ‘Exosome Specificity Factor’ (ESF) was theoretically hypothesized (4,33), its existence and functionality remained elusive. Only recently, Nrd1p was demonstrated to facilitate the exosomal degradation of Rho-induced transcription-elongation processing defective (76) mRNPs in *Saccharomyces* cerevisiae (46,47). Although these study unraveled a key role of Nrd1p in the degradation of Rho-induced faulty messages, Nrd1p/NNS was neither demonstrated as the primary recruiter of Rrp6 nor its contribution in the degradation of other kinds of aberrant nuclear mRNAs was addressed.

Physical interaction between Nrd1p and the cap-binding component Cbc1p, (a major component of CTEXT) and exosome component Rrp6p (33) inspired us to address the functional role of NNS complex in the nuclear mRNA decay. This inspiration was further bolstered when TRAMP4 component Trf4p was reported to interact with Nrd1p (77). As presented above, our results revealed that Nrd1p (and possibly NNS complex) participates in the degradation of all sorts of faulty mRNPs produced at different phases of mRNP maturation and the recruitment of Nrd1p (and possibly NNS) onto specific messages is extremely vital for their decay. While RNAPII plays an active role in the co-transcriptional recruitment of Nrd1p during transcription elongation, splicing, and 3’-end processing events via the RNAPII-CTD-Ser5-Nrd1p-CID interaction, its recruitment onto the export inefficient message is mediated by Pcf11p, via RNAPII-CTD-Ser2-Pcf11p-CID interaction (Table 4, Figs 7 and 8). This finding thereby implied that in the terminal phase of mRNP biogenesis, Nrd1p-RNAPII is displaced by Pcf11p-RNAPII once the RNAPII is phosphorylated at Ser2 (Fig 8). Substitution at the RNAPII-CTD is then followed by the recruitment of Nrd1p (NNS complex) on the aberrant mRNAs by Pcf11p (Fig 8). Alternatively, Pcf11p may directly recruit Nrd1p post-transcriptionally onto the export-defective messages without any involvement of RNAPII. Although it is not clear why Pcf11p is required for the recruitment of Nrd1p onto the export-defective message, it is possible that these messages may not have a strong association with RNAPII as they perhaps originate after the cleavage/polyadenylation reaction. At this stage, they appear to lose a strong association with RNAPII, while still remain attached to the transcription foci. This loose association necessitates the involvement of Pcf11p for their identification as aberrant RNA via the recruitment of Nrd1p. Future work would delineate this apparent mystery.

**Table 4:**
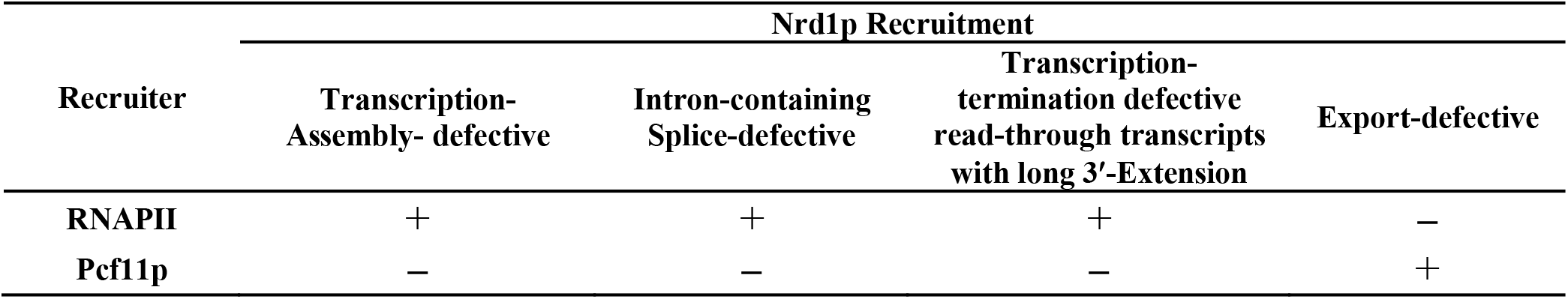
Recruitment profiles of Nrd1p on various aberrant nuclear mRNA targets.

Nevertheless, our data hints at an exciting possibility that the NNS complex by acting as an ESF co-ordinates the recruitment of either TRAMP or CTEXT differentially onto their distinct subsets of faulty messages (Figs 7 and 8). Although the underlying mechanism is still elusive, Nrd1p recruitment via RNAPII-Ser5-CTD-Nrd1p-CID interaction appears very critical for the selective recruitment of TRAMP (to target transcription elongation- and splice-defective messages). In contrast, its recruitment via RNAPII-Ser2-CTD-Pcf11p-CID interaction seems vital for the recruitment of CTEXT (to target and degrade export defective messages) (Fig 8). Indeed this conclusion explains our finding that the nuclear decay of the aberrantly long 3’-extended read-through transcripts requires the combined activity of TRAMP and CTEXT. These 3’-extended faulty transcripts are generated during the intermediate phase of mRNP biogenesis when the predominant population of RNAPII-CTD remains hyperphosphorylated consisting of both Ser5/Ser2 marks (78–80). Since, both Nrd1p and Pcf11p may remain associated with the hyperphosphorylated RNAPII-CTD having both Ser5-P/Ser2-P mark (68,81–83), this situation subsequently recruits both TRAMP (Recruited directly by Nrd1p) and CTEXT (Recruited indirectly by Pcf11p) onto these read-through transcripts presumably in the long 3’-extended segments (Fig 8). Our observation thus supports this conclusion that recruitment of Nrd1p onto these read-through messages in *nrd1*Δ^CID^ was not as low (about 40-50% compared to *NRD1*^+^ strain, Fig 7F) as those found in the cases of transcription elongation-defective and splice-defective messages (about 5-20% compared to *NRD1*^+^ strain, Fig 7D and E). This data thus strongly implied that in the *nrd1*Δ^CID^ yeast strain, the remaining 40-50% of the recruited Nrd1p onto these 3’-extended faulty messages was carried out via Pcf11p dependent manner.

Mining and re-analysis of previously published genome-wide high resolution Nrd1p and Nab3p binding map obtained from PAR-CLIP procedure (22,73,74) revealed that, while by and large the yeast mRNAome lacks the Nrd1p and Nab3p sites, several mRNAs, such as *NRD1, URA2, URA8, PCF11, FKS2* are enriched in the Nrd1p/Nab3p binding sites, whose expression were known to be regulated by Nrd1p (16, 18. 47–49). Interestingly, susceptibility of the export-defective special messages, *IMP3, NCW2* and *SKS1* to the Nrd1p-Nab3p-Sen1p complex even under normal condition in wild type cell prompted us to test if any of these mRNAs display any Nrd1p/Nab3p binding *in vivo*. As shown in Fig 9, analysis of the PAR-CLIP data showed that *NCW2* and *IMP3* harbor Nrd1p/Nab3p motifs and display strong Nrd1p/Nab3p binding *in vivo* as demonstrated by an overlapping T→C transition (Fig 9). This finding strongly corroborates and supports the idea that Nrd1p/Nab3p, following their binding to these mRNA targets, further co-ordinates the recruitment of the exosome/Rrp6p to facilitate their degradation. It should be noted here that Nrd1p/Nab3p binding to the native functional mRNA targets (such to *NRD1, PCF11, IMP3, NCW2* etc.) in a wild type cell is perhaps different in nature than those binding to aberrant mRNAs. In all our experiments presented above with specific mRNA biogenesis mutants, majority (if not all) of the mRNAs those are produced at conditional temperature of 37°C are aberrant in nature as characterized earlier (52, 54–59, 63). Interestingly, a thorough search for the putative Nrd1p/Nab3p binding motifs in the model transcription-elongation-defective (*ACT1* and *CYC1*), splice-defective (pre-*ACT1* and *pre-CYH2*) and 3’-end processing defective (*LYS2* and *CYC1*) mRNAs did not yield any well-defined motif (Fig 9A-B). However, the co-transcriptional recruitment profiles of some of these mRNAs strongly suggest that indeed Nrd1p preferentially binds to them when they are derived as aberrant/faulty messages despite lacking a defined motif. Thus, we favor the view that nature of the Nrd1p/Nab3p binding to the global or specific aberrant mRNAs is fundamentally different from their binding to the sites harboring the defined motif such as snoRNAs, snRNAs, CUTs, and the Nrd1p-target messages (*NRD1, URA2, URA8*, etc.). Consistent with this idea, recently Moreau et al (2019) also demonstrated that upon induction of the bacterial Rho-protein that triggers the expression of a vast majority of aberrant mRNAs in a transcriptome-wide scale, majority of the cellular Nrd1/Nab3p redistribute to the genomic loci of those protein-coding genes, which was sequestered in the genomic loci harboring snoRNAs and snRNAs before induction of the Rho-protein (47), many of them normally do not harbor any defined Nrd1p/Nab3p motif. Future research directed to determine the binding sites on these aberrant messages produced under conditional temperature of 37C would resolve this issue.

Our data imply that NNS complex qualifies well as a universal ESF by virtue of (i) exhibiting a uniform affinity to all categories of aberrant nuclear messages, and (ii) having an ability to identify and delineate a distinct set of aberrant message from a massive pool of cellular mRNAs via its increased occupancy of itself on these messages. This conclusion is strongly supported by the increased co-transcriptional recruitment observed onto the aberrant mRNA targets generated in THO mutant strains (e.g., mRNAs from *hpr1*-Δ at 37°C compared to 25°C, see Fig 7A and B) or onto the special messages (*NCW2* vs. *CYC1*, see Fig 7C). Even though the molecular basis of the distinction between the normal/functional and aberrant/faulty messages is not clear at this point, it appears from this work that this difference may be accomplished co-transcriptionally by the presence of Nrd1p which is selectively recruited more on the aberrant messages (Fig 7A-C). It is reasonable to assume that during transcription elongation, a competition exists between the binding of the mRNP biogenesis factors (such as THO/splicing/export factors) and the association of NNS with the transcribing messages. In case of a normal message, the mRNP biogenesis factors are promptly deposited on the maturing message thereby excluding the binding of the Nrd1p. During the transcription elongation of the aberrant messages, in contrast, binding of processing factors becomes impaired, leading to retardation of the speed of transcription allowing more time for NNS to bind co-transcriptionally to these aberrant messages. However, it is currently unknown whether RNAPII actively coordinates the selective recruitment of Nrd1p onto specific types of aberrant messages during various phases of transcription elongation and mRNP maturation events. Alternatively, targeted Nrd1p recruitment onto aberrant messages may also result from the competition between the mRNP biogenesis factors and NNS complex, as described above. Nevertheless, this mystery leaves us with an opportunity to investigate further, in an opportunity to gain an insight into the distinguishing feature(s) of aberrant messages that leads to the subsequent recruitment of Nrd1p (possibly NNS complex) to facilitate their degradation. The aligned reads were then investigated for T → C transitions using Integrative Genomics Viewer (REF.).

## Materials and methods

### Nomenclature, strains, media, and yeast genetics

Standard genetic nomenclature is used to designate wild-type alleles (e.g., *ACT1, CYC1, CYH2, LYS2, HPR1*), recessive mutant alleles (e.g., *lys2-187, nup116*-Δ, *nrd1-1, nrd1-2, nrd1*Δ^CID^) and disruptants or deletions (e.g., *cbc1::TRP1, cbc1*-Δ, *rrp6::TRP1, rrp6*-Δ). An example of denoting a protein encoded by a gene is as follows: Nrd1p encoded by *NRD1*. The genotypes of *S. cerevisiae* strains used in this study are listed in **Table S1**. Standard YPD, YPG, SC-Lys (lysine omission), and other omission media were used for testing and growth of yeast (84). Yeast genetic analysis was carried out by standard procedures (84).

### Plasmids and oligonucleotides

The plasmids were either procured from elsewhere or were constructed in this laboratory using standard procedures. All the oligonucleotides were obtained commercially from Integrated DNA Technology. The plasmids and oligonucleotides used in this study are listed in **Tables S2 and S3**, respectively.

### Plate Assay for suppressibility of the *lys2-187* mutation

A series of 10^-1^ dilution of suspension of 10^5^ cells per ml were spotted on YPD, and lysine omission medium (SC-Lys) and the plates were incubated at 30°C for either 3 (for YPD) or 4-5 days (for Sc-Lys) medium followed by capturing the image of the cell growth as described previously (15).

### RNA analyses and determination of steady-state and decay rate of mRNAs

Total RNA was isolated as described earlier (16) by harvesting appropriate yeast strains followed by extracting the cell suspension in the presence of phenol-chloroform-IAA (25:24:1) and glass bead. Following the extraction, the RNA was recovered by precipitation with RNAase-free ethanol. For the preparation of cDNA, total RNA was treated with DNase I (Fermentas Inc., Pittsburgh, PA, USA) to remove genomic DNA at 37°C for 30 minutes. The reaction was stopped by adding 1μl of EDTA and incubating at 65°C for 10 minutes, followed by first-strand cDNA synthesis using Superscript Reverse Transcriptase (Invitrogen) using Random Primer (Bioline Inc.) by incubating the reaction mixture at 50°C for 30 minutes. Real-time qPCR analyses were performed with 2–3 ng of cDNA samples for *ACT1, CYC1, LYS2, NCW2, SKS1*, etc. and 30 ng for intron-containing splice defective messages, *ACT1* and *CYH2* in *prp2-1* strains to determine the steady-state levels as described previously (16) decay rate of a specific mRNA was determined by the inhibition of global transcription with transcription inhibitor 1, 10-phenanthroline (Sigma-Aldrich) at 25°C or 37°C (as mentioned) and is described previously (16). Briefly, the specific strain was grown at 25°C till mid-logarithmic phase (special messages), or an additional step was added for temperature-sensitive mutant (*hpr1*-Δ, *prp2-1, rna14-1*) of shifting the culture from 25°C to 37°C for 2 hours. This was followed by the addition of 1, 10-Phenanthroline to the growing culture at a final concentration of 100 μg/mL and a shift in the withdrawal of a 25 ml of aliquots of culture at various times after transcription shut off. Messenger RNA levels were quantified from cDNA by real-time PCR analysis, and the signals associated with the specific messages were normalized against *SCR1* signals. The decay rates and half-lives of specific mRNAs were estimated with the regression analysis program (Graphpad Prism version 7.04) using a single exponential decay formula (assuming mRNA decay follows first-order kinetics), y = 100e^-bx^ was used.

### Real-Time PCR

Following the synthesis from total RNA samples, each cDNA was first quantified using Qubit^®^ds DNA HS Assay Kit (Life Technologies, USA) following their recommendation. 2 to 3 ng of quantified cDNA was used to quantify the levels of specific mRNAs such as *CYC1*, *ACT1, IMP3, NCW2*, and *LYS2*, etc. by qPCR assays by using target-specific primers and standard SYBR Green Technology using Power SYBR^®^ Green PCR Master Mix (Applied Biosystems). qPCR assays were run in triplicate and conducted using an Applied Biosystems StepOne^TM^ real-time PCR system to determine the absolute quantification of individual target mRNAs. For each target either ΔΔ^-Ct^ method was used, or purified DNA templates were prepared by PCR, followed by Gel purification and diluted serially to generate standard curves from 10^2^ to 10^9^copy number/reaction versus threshold cycle (Ct), determined from fluorescence by the StepOne^™^ software version 2.2.2 (Applied Biosystems). Standard curves were used to determine the amplification efficiencies (E =10^(−1/n)^, where n = slope of the standard curve) in SYBR Green assays, as well as copy numbers of templates in experimental samples.

### Protein analyses by western blot

Total protein was isolated from specific yeast strains (WT and *nrd1*Δ^CID^) grown overnight at 30°C in YPD broth. Following centrifugation at 5000 rpm for 7 min the cell pellets were quickly frozen in liquid nitrogen and stored at −70°C. Frozen pellets were thawed on ice and resuspended in 1 ml of Buffer A (50 mM Tris–HCl pH 7.5, 150 mM NaCl, 5 mM EDTA, 1 mM DTT, 1 mM PMSF) supplemented with Protease Inhibitor Cocktail (Abcam ab201111, UK) and the cells were lysed by vortexing 10–15 times with glass beads followed by clarification of the particulate fraction by centrifugation. Supernatants were collected by centrifugation at 11 000 rpm for 20 min and saved as the total soluble protein fraction for further analysis. Protein concentration was determined by Bradford reagent assay kit (Bio-Rad Inc., Valencia, CA, USA). For Western Analysis, 60 μg of total protein was used, which was resolved in 10% SDS polyacrylamide gel. The separated proteins were transferred to PVDF membrane at 50-100mA O/N for immunobloting. Blots were blocked with 5% skimmed milk in Tris-buffered saline (10mMTris, 150mMNaCl, 0.1% Tween 20) and incubated with primary antibodies for specific proteins for 1 h at room temperature in following dilutions in 5% BSA: a) Rabbit Polyclonal anti-Nrd1 (1:1000), and b) Mouse Monoclonal anti-Tubulin (1:2000). Blots were further washed in 1X TBS with 0.1% Tween 20 (TBST) and incubated in HRP-conjugated secondary anti-rabbit or anti-mouse antibody each diluted at 1:3000 in wash buffer for 1 h at room temperature. Immunoreactive bands were developed and detected by chemiluminescence (ECL imager kit, Abcam) and the images were captured by X-ray film or myECL Chemidoc Imager (Thermo Scientific, USA).

### Chromatin Immunoprecipitation (ChIP) Assay

Chromatin preparation was essentially performed following the procedure described earlier (85). The *NRD1*^+^ and *nrd1*Δ^CID^ strains or *PCF11*^+^ and *pcf11-2* strains in the different backgrounds were used for this study. Two fifty milliliters of cells grown till OD_600_ ≈ 0.5 (≈ 10^7^cells/ml) and were fixed with 1% formaldehyde for 20 min. The reaction was stopped by the addition of glycine, and the cells were washed and lysed with glass beads to isolate chromatin. The cross-linked chromatin was sheared by sonication to reduce average fragment size to ≈500bp. Chromatin-IP was carried using the ChIP assay kit (EZ-ChIP ^TM^; Cat#17-295) from Merck Millipore. Immunoprecipitation of 120 μg Chromatin fractions (≈ 100μl) from each strain was performed using anti-Nrd1p antibody incubated overnight at 4°C. Incubation with Protein G agarose bead was followed by washing and chromatin elution, the eluted supernatants and the input controls were incubated with Proteinase K for 1h at 37°C followed by 5h at 65°C to reverse cross-link the precipitated DNA from the complexes. DNA was purified using the DNA-clean up column provided with the kit. The immunoprecipitated DNAs (output) were quantified by real-time PCR (as mentioned above) using three sets of primers located along a specific genes coding sequence (5’-end, middle, and 3’-end) and normalized to a half dilution of input DNA. Amplifications were done in duplicate for each sample (technical replicate), averages, and standard deviations were calculated based on three independent experiments (biological replicate).

### Mining and Analysis of The previously deposited PAR-CLIP data

The processed wig files for the Nrd1 Arc Lamp Crosslinking dataset were downloaded from Gene Expression Omnibus (86) found under series number GSE31764 (Originally deposited by Creamer et. al. 2011) (73) The files were loaded into Integrative Genomics Viewer (87) using which, the Nrd1 PAR-CLIP binding plots were obtained. In order to find the sites of T → C transition, the raw reads were downloaded from SRA (88) and aligned to the *Saccharomyces cerevisiae* reference genome version R64-1-1, downloaded from https://asia.ensembl.org/ using RNA STAR(89), on the European public server of the galaxy project (https://usegalaxy.eu/)(90). The aligned reads were then investigated for T → C transitions using Integrative Genomics Viewer (87).

**Figure 10.**
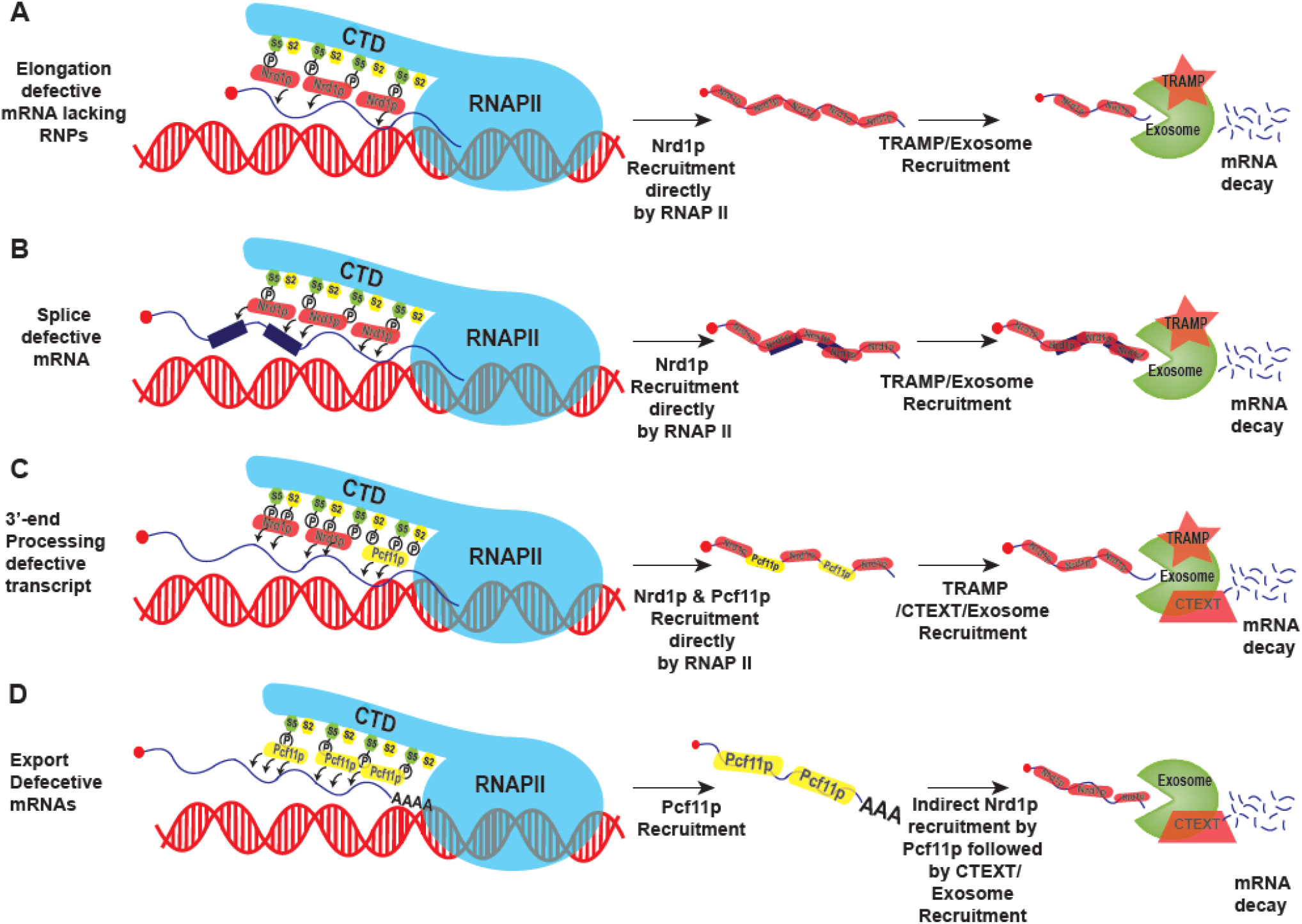
Model elucidating the Nrd1p-dependent recognition, and degradation of transcription-assembly-defective (A), splice-defective (B), 3’-end processing defective (C), and export-defective (D) messages in baker’s yeast. During transcription initiation, elongation, and termination steps, each message is monitored for the existence of aberrancies, probably by RNAP II. Nrd1p is recruited either directly (A-B) onto the transcription-assembly- and splice-defective messages during the initial stages of transcription/mRNP biogenesis (RNAPII-CTD predominantly consists of Ser-5 marks) leading to the recruitment of the TRAMP/exosome, or indirectly via Pcf11p (D) onto the export-defective messages during the late stage of mRNP biogenesis (RNAPII-CTD predominantly consists of Ser-2 marks) leading to the recruitment of CTEXT/exosome. During the intermediate phase of mRNP biogenesis, (RNAPII-CTD predominantly carries Ser-5-2 marks), Nrd1p is recruited both directly by RNAPII-CTD as well as indirectly by Pcf11p, leading to the recruitment of TRAMP/CTEXT/exosome. In every case, the recruitment of exosome onto the aberrant messages leads to their nuclear degradation. In (B), the tiny blue boxes depict the introns.

## Abbreviations

TRAMP: **<u>TR</u>**f4-**<u>A</u>**ir1/2-**<u>M</u>**tr4 **<u>P</u>**olyadenylation
CTEXT: **<u>C</u>**bc1-**<u>T</u>**if4631-dependent **<u>EX</u>**osomal **<u>T</u>**argeting
NNS: **<u>N</u>**rd1p-**<u>N</u>**ab3p-**<u>S</u>**en1p

## Acknowledgments

We gratefully acknowledge Dr. Stephen Buratowski (Department of Biological Chemistry and Molecular Pharmacology, Harvard Medical School, Boston, MA, USA) for kindly sharing *nrd1-1, nrd1-2*, *nrd1*Δ^CID^, *sen1-1*mutant strains, anti-Nrd1antibody, and various *NRD1* plasmids. We also acknowledge Dr. Maurice Swanson (University of Florida, Gainesville, FL, USA), Dr. Jeffry Corden (Johns Hopkins School of Medicine, Baltimore, MD, USA) for *nab3-10* strain, Dr. Athar Ansari (Wayne State University, Michigan, USA) for the *pcf11-2* mutant yeast strains, and Dr. Scott Butler for providing us with the Anti-Rrp6p antibody (University of Rochester, Rochester, NY, USA). We thank Dr. Scott Butler, Dr. Satarupa Das, Dr. Arindam Chakraborty, and the members of the Das Laboratory for critically reading this manuscript. We also thank the anonymous reviewers for their critical comments and constructive suggestions, which certainly enriched this manuscript.

## Supplementary Tables

**Table S1:**
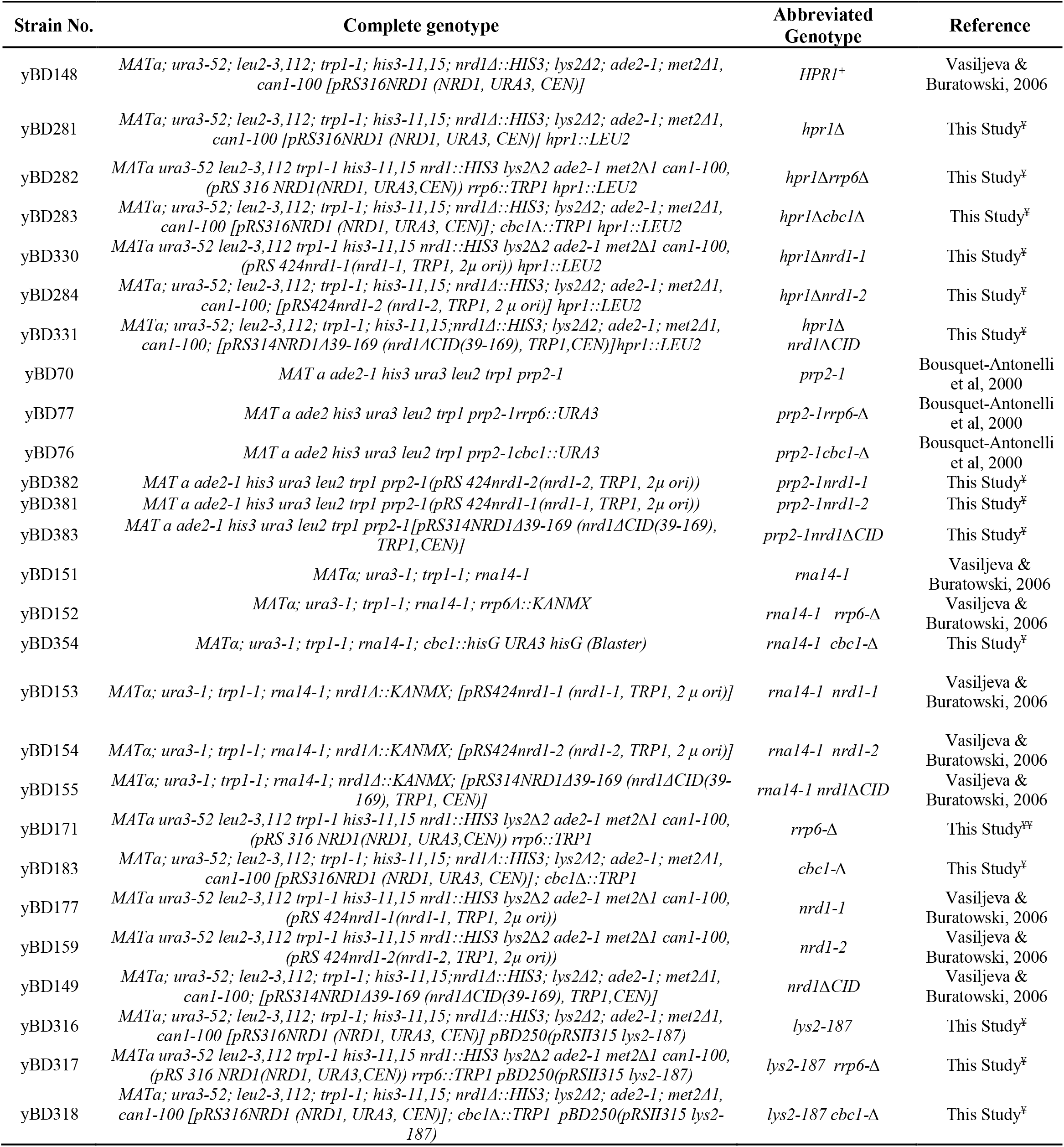
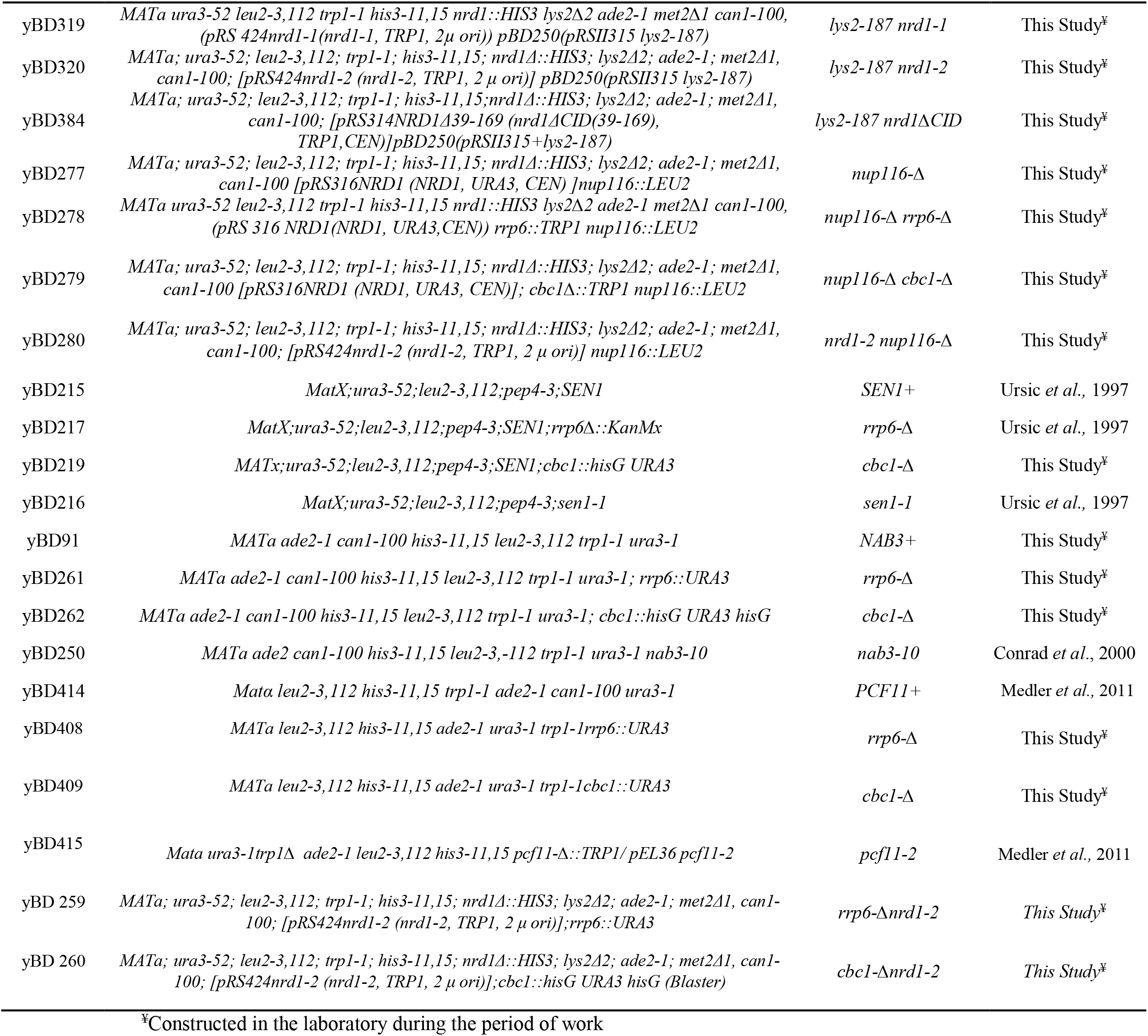
List and Genotypes of Yeast Strains used in this study.

**Table S2:**
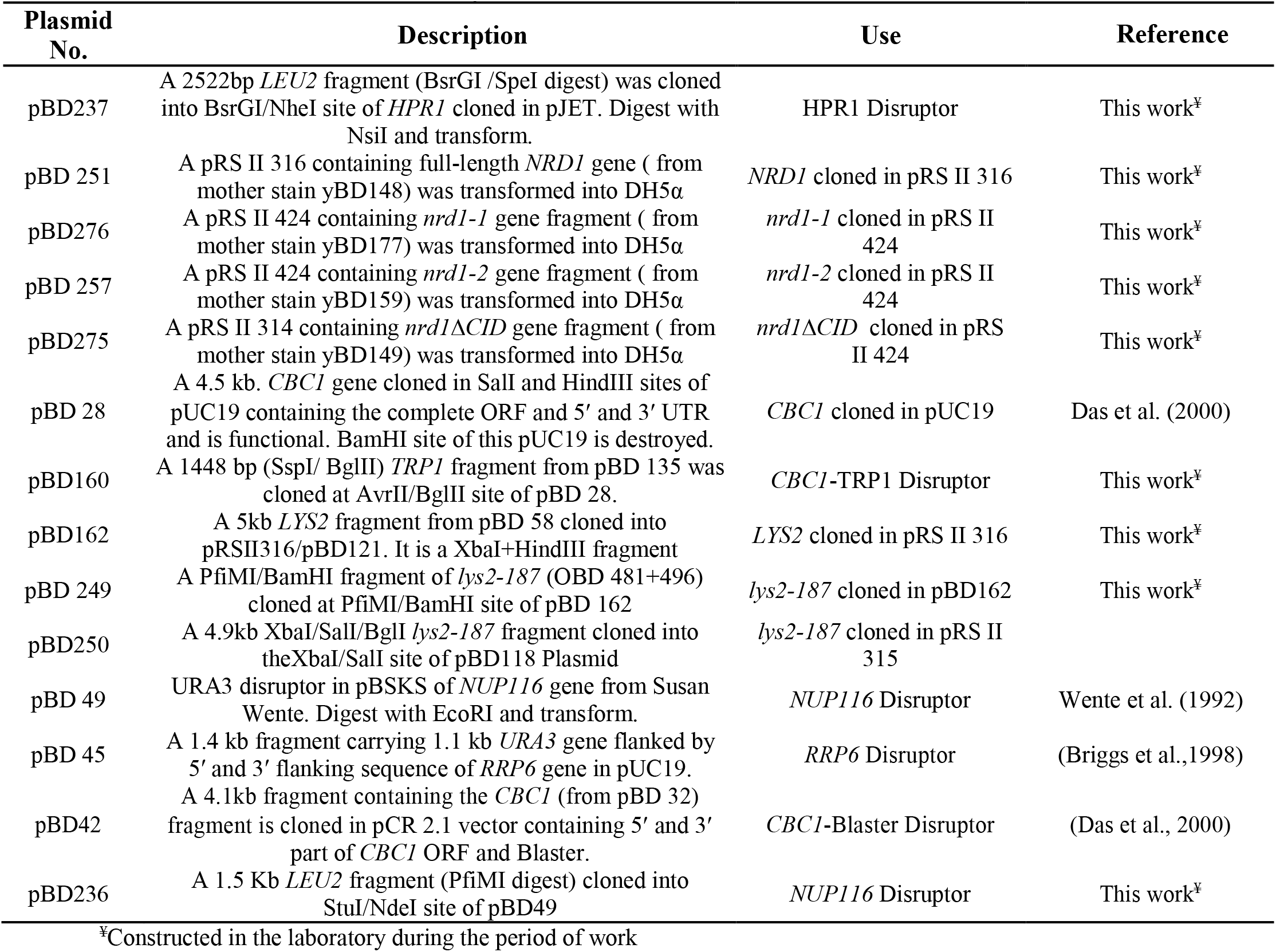
List of Plasmids used in this study.

**Table S3:**
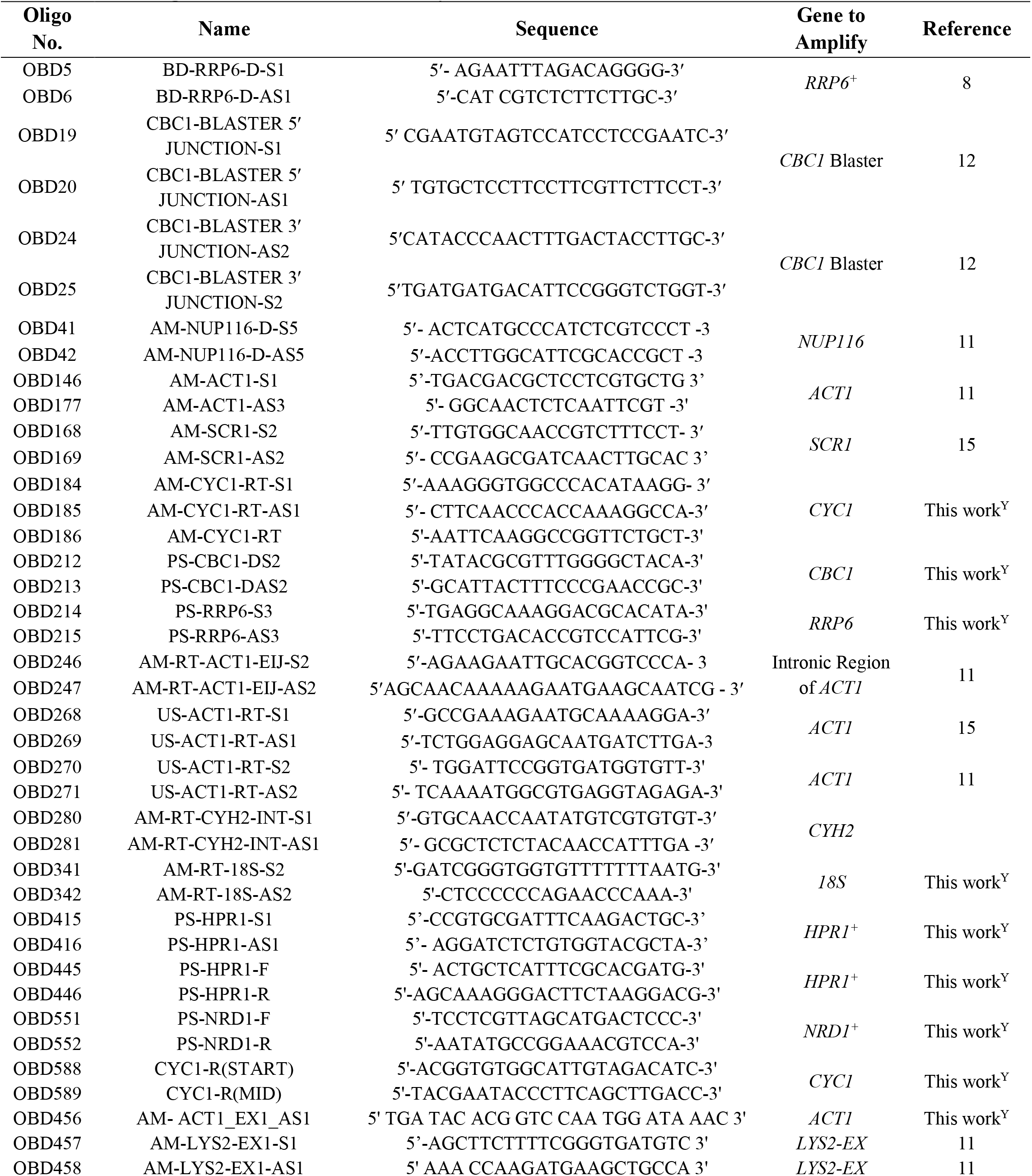
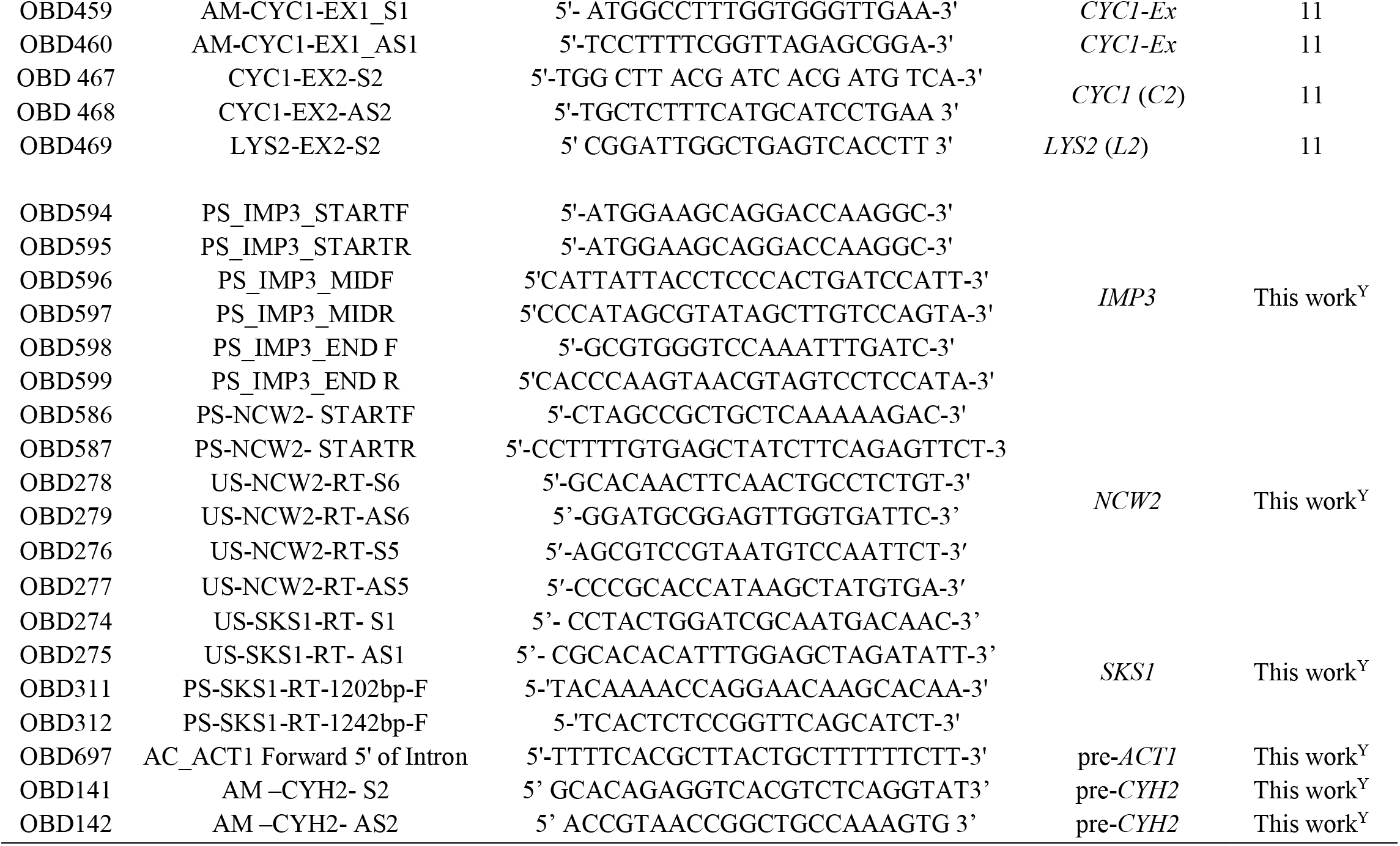
List of Oligonucleotides used in this study.

## Legends to the Supplementary Figures

**Figure S1.**
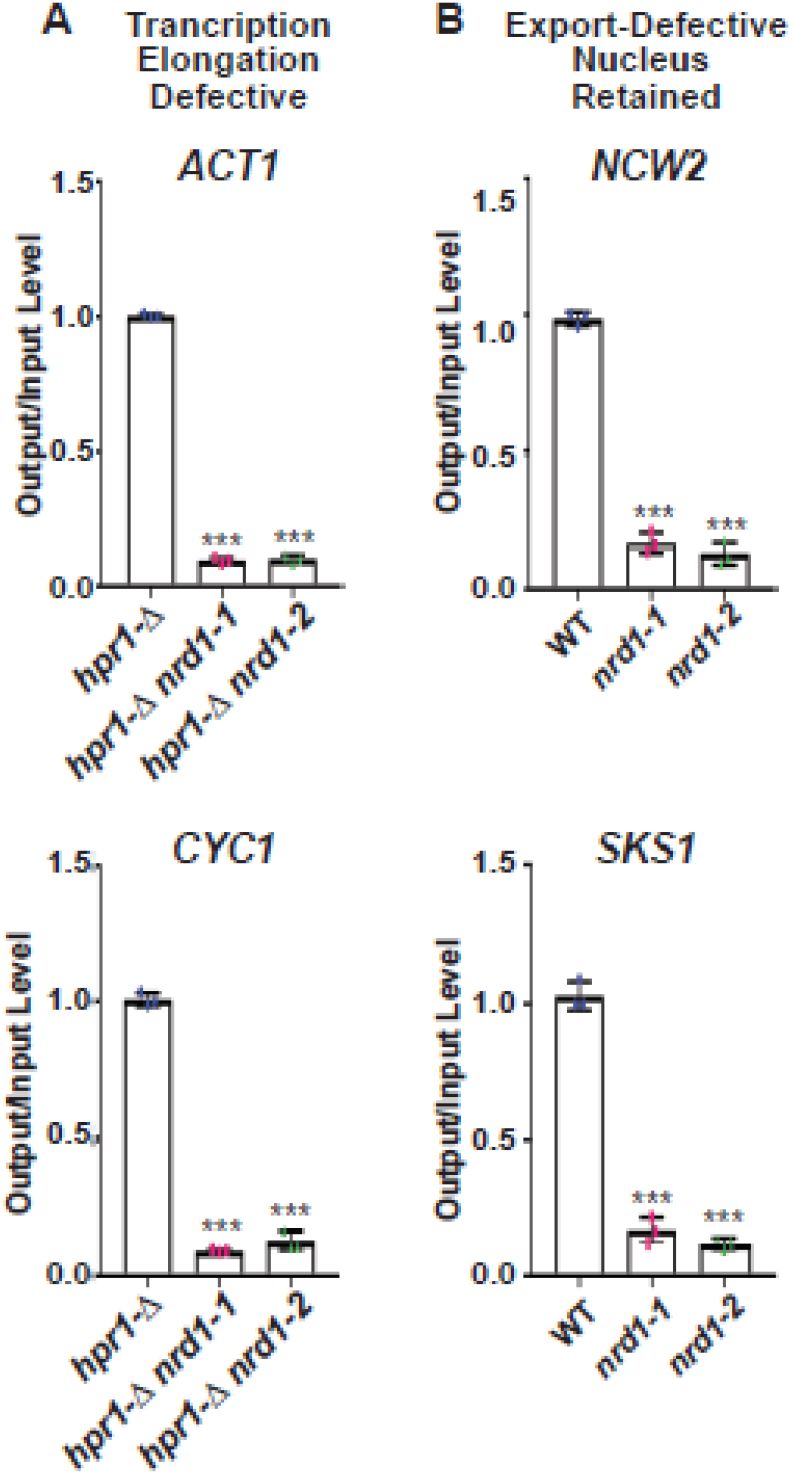
Co-transcriptional recruitment of full-length Nrd1p and truncated proteins in wiild-type and *nrd1-1* and *nrd1-2* strains for (A) transcription assembly-defective, *ACT1*, and *CYC1* mRNA and (B) export-defective special messages, *NCW2* and *SKS1* mRNAs. All immunoprecipitation using specific Nrd1p antibody followed by the recovery of the precipitated DNA and qPCR analyses was performed exactly as described in materials and methods and the legend of Figure 6. The primer sets specific for the central region of corresponding to the ORF of each target mRNA were used for the qPCR analyses to generate these scatter plots. P-values estimated from Student’s two-tailed t-tests for a given pair of test strains for each message, is presented with the following symbols, * <0.05, **<0.005 and ***<0.001, NS, not significant.

**Figure S2:**
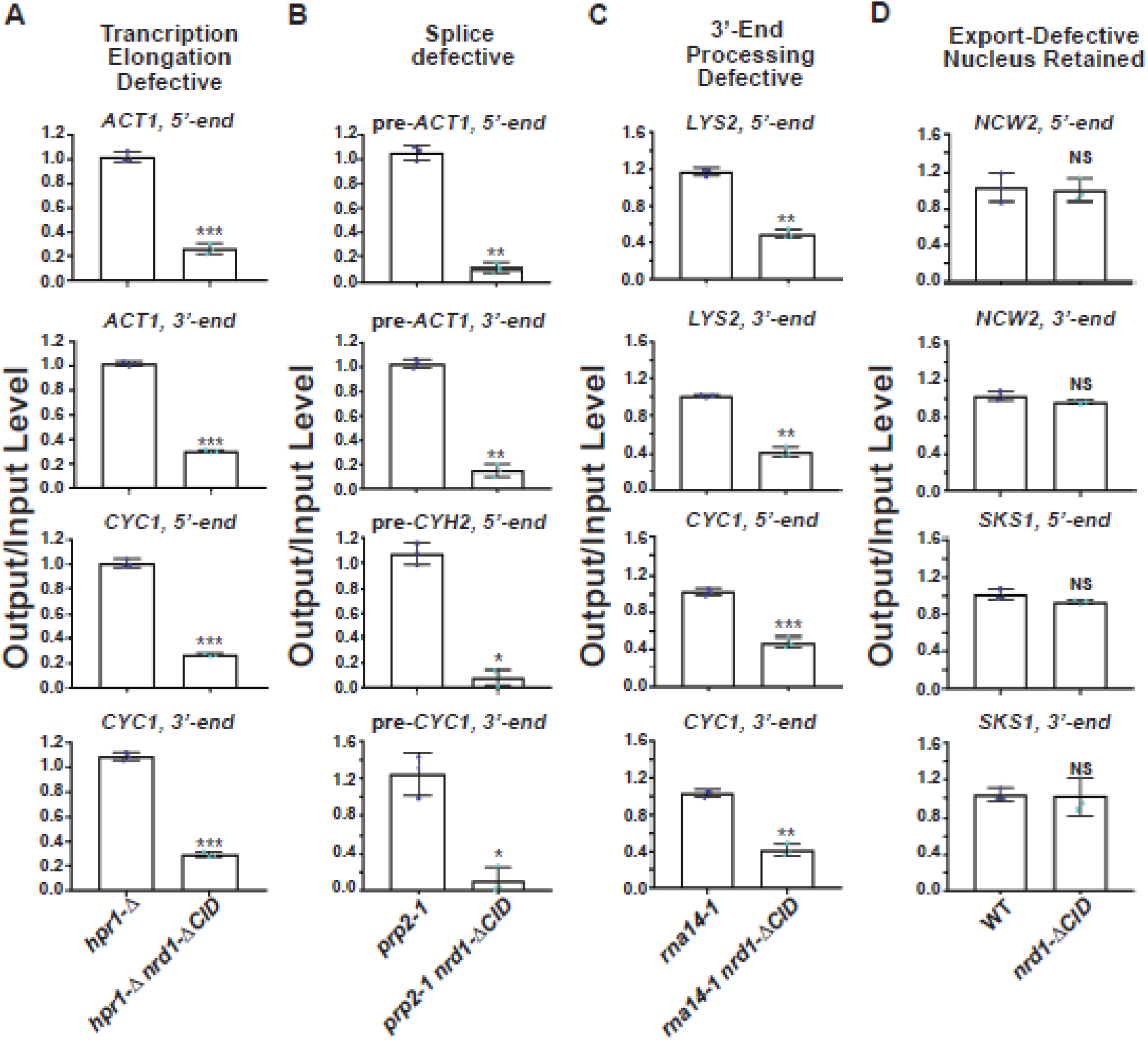
Co-transcriptional recruitment of full-length Nrd1p and Nrd1p lacking the *CID* domain on (A) transcription assembly-defective, (B) splice-defective and (C) 3’-extended aberrant messages, and (D) export-defective special messages. Occupancy profile of full-length Nrd1p and Nrd1p lacking its *CID* domain of the (A) transcription-assembly-defective *ACT1* and *CYC1* mRNAs at 37°C in isogenic *hpr1*-Δ and *hpr1*-Δ *nrd1*Δ*CID* strains, (B) splice-defective intron-containing pre-*ACT1* and *pre-CYH2* at 37°C in isogeneic*prp2-1* and *prp2-1 nrd1*Δ*CID* strains (C) 3’-extended read-through *CYC1* and *LYS2* transcripts at 37°C in *rna14-1* and *rna14-1 nrd1*Δ*CID* strains and (D) nucleus-arrested special messages *NCW2* and *SKS1* at 25°C in wild-type (*NRD1*^+^) and *nrd1*Δ*CID* strains. All immunoprecipitation using specific Nrd1p antibody followed by the recovery of the precipitated DNA and qPCR analyses was performed exactly as described in materials and methods and the legend of Figure 6. The primer sets specific for the 5’-end and 3’-end corresponding to the ORF of each target mRNA were used for the qPCR analyses to generate these scatter plots. P-values estimated from Student’s two-tailed t-tests for a given pair of test strains for each message, is presented with the following symbols, * <0.05, **<0.005 and ***<0.001, NS, not significant.

